# Design of an ultrafast pulsed ponderomotive phase plate for cryo-electron tomography

**DOI:** 10.1101/2022.05.09.491239

**Authors:** Daniel X. Du, Anthony W. P. Fitzpatrick

## Abstract

Ponderomotive phase plates have shown temporally consistent phase contrast is possible within electron microscopes *via* high fluence static laser modes resonating in Fabry-Perot cavities. Here, we explore using pulsed laser beams as an alternative method of generating high fluences. We find through forward-stepping finite element models that picosecond-or-less interactions are required for meaningful fluences phase shifts, with higher pulse energies and smaller beam waists leading to the predicted higher fluences. An additional model based on quasiclassical assumptions is used to discover the shape of the phase plate by incorporating the oscillatory nature of the electric field. From these results, we find the transient nature of the laser pulses removes the influence of Kapitza-Dirac diffraction patterns that appear in the static resonator cases. The addition of a second laser aligned 90° to the first induces anisotropy to the shape of the phase plate. By incorporating a shifting-electron-beam algorithm, the effects of a finite electron beam crossover are also simulated. A total pulse energy of 8.7 μJ is enough to induce the required π/2 phase shift for Zernike-like phase microscopy. As a brief thought experiment, we also explore the usage of high frequency lasers in a standard electron emission scheme to see if a pulsed electron beam is even necessary. Ultimately, frequency requirements limit the laser to nanosecond pulse durations, causing the required pulse energies to reach unreasonable levels before adequate phase shifts are achieved.

## Introduction

The imaging of biological specimens, and soft matter in general, has significantly advanced to the point where high-resolution structure determination of protein complexes is becoming increasingly commonplace (1–4). However, this process itself is not without its difficulties, both for single-particle cryo-electron microscopy (cryo-EM) and cryo-electron tomography (cryo-ET). At resolutions better than 4 Å, required to identify side chains, tens of thousands of particles are acquired, aligned, and superimposed to produce sufficient contrast for structure determination (5–7). In addition, numerous software-based contrast enhancement techniques must be applied to aid in segmentation and annotation of cryo-electron tomograms (5, 8–14). Much of the reason for the time and effort invested in these forms of image-processing can be attributed to the weak scattering of electrons from soft/low-atomic number (low-*Z*) matter (15–17).

Indeed, this problem has originated in optical microscopy and was eventually mitigated either by an intentional defocusing of the objective to acquire phase-dependent fringes or through the development of the Zernike phase microscope. The Zernike phase microscope enhances phase contrast by altering the phase of the unscattered light transmitted from a specimen by a factor of *π*/2. This additional factor creates destructive interference, much like Bragg diffraction, and thus enhances contrast at feature boundaries (18). This concept has long been theoretically extended to the electron microscope, but only recent advancements have made such phase shifts possible to an electron beam (19–23). Most prominently, the Volta phase plate and other such material-based phase plates have demonstrated sharp increases in contrast, though these methods still struggle with electrostatic charging and beam damage (20, 23–28).

An alternative method proposed by Reed in a patent used a ponderomotive field potential generated by focused laser beams in a photon-induced near-field electron microscopy (PINEM)-like effect to alter the phase of electrons in a narrow field of view, with the aim of avoiding any deleterious effects such as material damage or charging (22, 29). Recently, this idea has been implemented in two continuous wave forms with the inclusion of resonators to reach the required powers for sufficient Compton scattering. The first involves a Fabry-Perot resonator while the second uses a ring microresonator. Already, early results show similar contrast enhancements to material phase plates and work continues to mitigate the aberrations that occur from scattering and physically changing the column of the TEM (30–33).

The usage of a continuous resonating beam enables the formation of a stable continuously replenishing phase plate, thereby bypassing the charging and damage concerns of material phase plates. However, a resonator is not the only method to localize the power in a laser beam. As an alternative, pulsed beams, capable of being produced *ex situ*, are well known to achieve high instantaneous fluences (34). Here, we explore the proposed pulsed ponderomotive phase plate, as demonstrated in Figure 1, and determine the various experimental parameters required for efficient phase contrast *via* a forward-stepping finite element model. It is our intent to understand if the sharp increase in fluence at the beam waist of an ultrafast laser is sufficient to generate the required *π*/2 phase shift relative to the unscattered electrons and to further explore the effects of changing electron and laser spatiotemporal pulse widths. There are two models we will use, one resulting from a fundamental scattering frame and one from a quasiclassical frame, which have been adapted to fit a forward-stepping algorithm (21, 30). The two models are used for distinct purposes that are discussed later. The results of these investigations are aimed at guiding future development in the pulsed ponderomotive phase plate.

**Figure 1.**
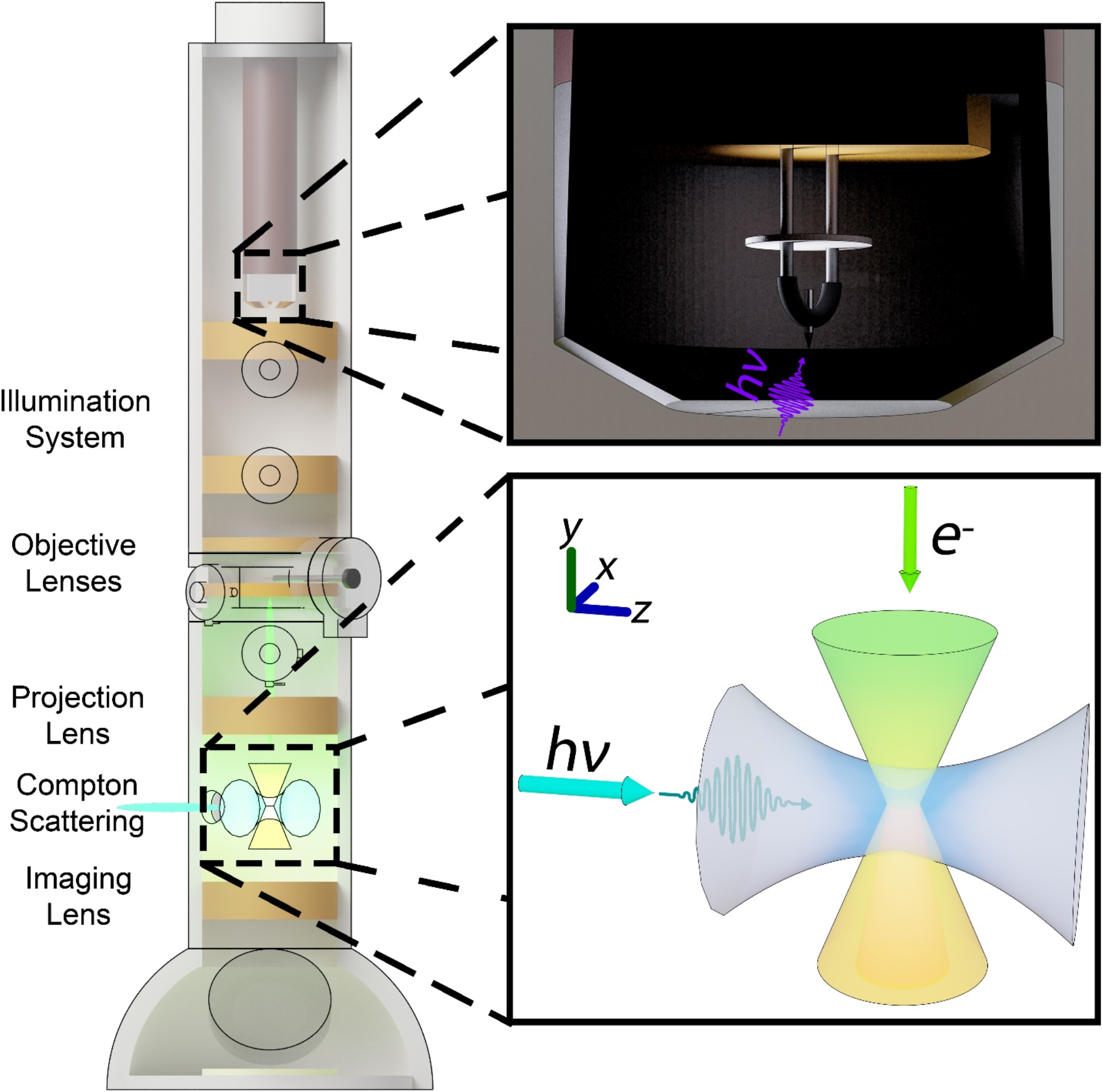
A cross-section of the pulsed ponderomotive phase plate operating inside an ultrafast electron microscope. Each of the major sections of the transmission electron microscope (TEM), including the region with the ponderomotive phase plate, are labeled. Here, the region with Compton scattering is specifically targeted to a location where the back focal plane of the specimen is reformed, allowing for the scattering against the direct (000) peak. In the upper inset, a schematic demonstrates the generation of the pulsed electron beam with an ultraviolet laser pulse. In the bottom inset, a closeup of the ponderomotive phase plate demonstrates where the laser beam waist is positioned such that it has the highest density of photons for Compton scattering. Likewise, the electron beam experiences a refocusing of the back focal plane at the exact same location. Here, only the direct beam is shown.

### Initial Experimental Bounds with the Scattering Method

The most prominent aspect of the pulsed ponderomotive phase plate is determining whether a pulsed laser beam even has enough photon density to generate the required *π*/2 phase shift for Zernike phase contrast. Here, we use one of the theories set forth by Müller *et. al*. and described by a four-particle Feynman diagram (21). The phase shift from the ponderomotive effect is then generally described by the following equation:

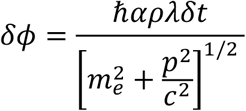

where *α* is the fine-structure constant, *ρ* is the instantaneous photon density in the presence of the scattered electron, *λ* is the wavelength of the photons, *p* is the momentum of the electron, and *δt* is the measured interaction time. Using this equation, we simulated the interaction between a photon pulse of 515 nm in wavelength and an electron pulse of 200 keV electrons. To keep the problem simple, the laser beams used in the simulation are of a TEM_00_ gaussian beam, adapted to fit an ultrafast temporal profile (35). The equation used to approximate an ultrafast gaussian pulse is described by:

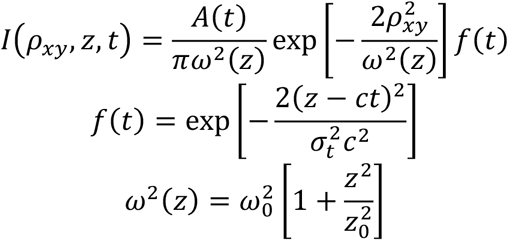

where *A*(*t*) is a time-varying parameter used to normalize the pulse to the pulse energy, *ω*(*z*) is the gaussian beam diameter, *z*_0_ is the Rayleigh range, *ρ_xy_* is the radial distance from the travel axis of the laser, *σ_t_* is the time resolution, *c* is the speed of light, and *z* is the distance down the travel axis. More on the implementation of this simulation is discussed in the methods section and in the Supplemental Information, Section 1 and Section 2. The results of these simulations are summarized in Figure 2. An example contour is provided as well, which describes the result of interactions from electrons traveling at different angles through the laser beam. For this section, each pixel of the contour represents an electron path of different slope that all pass through the center of the laser beam (marked by the direction of electron travel) in a cone as the pulse center of the laser beam passes through the origin. As such, electrons represented by pixels on the right travel counter to the direction of the laser beam in the *xz*-plane. Electrons represented by pixels on the left travel in the same direction as the laser beam in the *xz*-plane. Electrons in pixels at the top travel towards -*x* while those at the bottom travel towards +*x*. As is the case in electron microscopes, these electrons also all primarily travel in the -*y* direction, which is orthogonal to the plane of the contour plot and extends into the face of the page.

**Figure 2.**
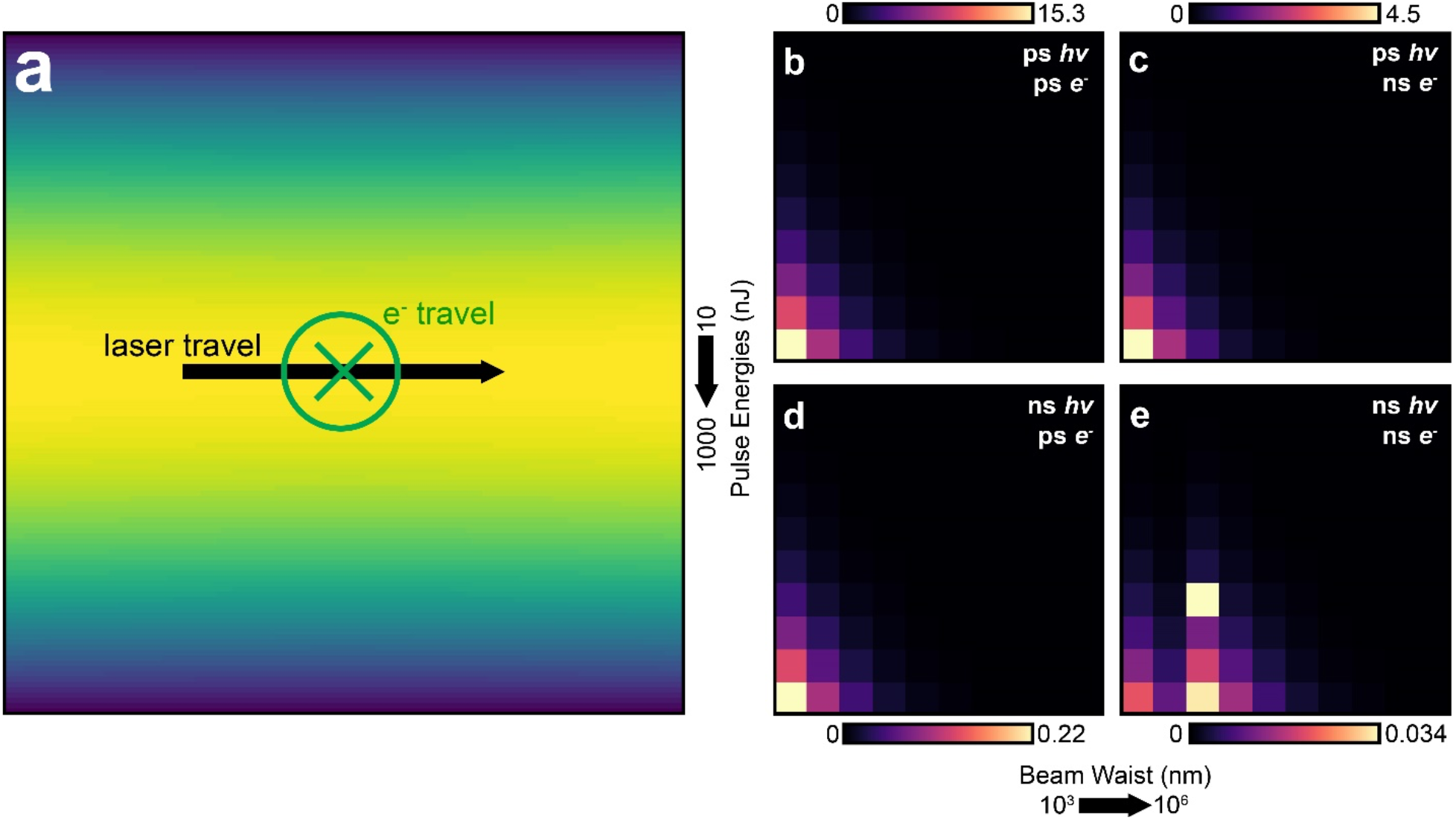
A demonstration of the simulated results alongside contour plots of the peak difference in phase shift between the point of maximum shift and the point of minimum shift. (a) A typical result from the simulations involving the scattering method, with the direction of laser travel progressing from left to right and the electron beam traveling into the page. (b-e) Contour plots of the phase shift difference in the electron beam. The pulse energies are along the *y*-axes, with values from 10 nJ to 1000 nJ in logarithmic form. The *x*-axes describe the beam waist size from 1 μm to 1 mm. Colorbars have been included with each contour plot showing the phase shift, with each colorbar normalized to the peak shift in the contour plot. The temporal resolutions (1σ) of both the laser and electron beams are shown in the top right.

The immediately noticeable result is the expected trend of higher beam powers generating a higher amount of phase shift, a result that arises directly from the interaction equation and the photon density term. Similarly, the photon density is increased when the beam waist narrows, another expected result from the gaussian pulse equations. Aside from the mathematical consequences to the phase shift resulting from the choice of model, the π/2 Zernike phase shift is exceeded in only two of the temporal resolution cases tested. Here, we have tested the model by varying the laser and electron pulses between 10 ps and 1 ns in pulse duration (*i.e*. standard deviation). Out to 1 μJ of pulse energy, the phase shift exceeds the π/2 requirement when the laser beam is condensed to 10 ps, as expected based on the resulting instantaneous photon densities. In the case of matching electron and laser pulse durations, the expected phase shift is 15.3 radians at the peak, while a mismatched nanosecond electron beam generates a 4.5 radian phase shift.

We note the odd behavior in the contour plots for the cases of ns laser and electron pulses. This behavior is a direct result of lack in time granularity in electron slice stepping, which is unfortunately necessary to alleviate computation overhead. Without such overhead, we would expect a monotonic increase towards bottom left (*i.e*. smaller beam waist, higher pulse energy). Future simulations may account for this by incorporating non-uniform meshes, though it is not a primary concern as we are more interested in the trend, order-of-magnitude phase shifts, and shape of the pulsed phase plate.

It is important to further discuss the reporting of phase as an average phase difference across the entirety of an electron pulse. Here, we note that while the average phase increase in the case of a 10 ps laser pulse and 1 ns electron pulse reaches the adequate average phase shift necessary for phase contrast, a deeper exploration of the total number of electrons that shift in phase reveals the shift only exists in the pulse duration where both the laser and electrons overlap. As such, only a limited section of the electrons in this scenario shows any appreciable phase shift. This is further explored in the Supplemental Information, Section 3. From the results of these preliminary simulations, we focus attention on the most promising cases of short-duration 10 ps pulses for much of the remainder of the text.

### Single- and Double-Laser Quasi-classical Approximation for Shape Determination

The shape of the phase plate is of concern, as it has a direct effect on the contrast transfer function (CTF) and analysis of the resulting image (36). It is well known that in the context of a continuous wave, the oscillatory nature of the photons form a Kapitza-Dirac diffraction grating which is also imprinted onto the phase shifted electrons (37). In the pulsed case, it is unclear if this diffraction grating would be as clearly resolved. We hypothesize that the transient nature of both the electron and laser pulses would smear the original Kapitza-Dirac diffraction grating and would approach the results from the scattering model. The central equation governing the shape of the phase plate resulting from scattering against the oscillatory fields is given by the following:

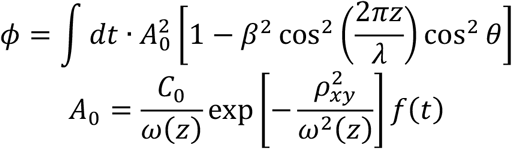

where *C*_0_ is an arbitrary vector amplitude measured in volts, *β* = *v*/*c*, and *θ* is the polarization of the laser (here set to zero). This model has been adapted from Turnbaugh *et. al*. and once again changed to fit ultrafast pulse interactions (30). Further information on this model is provided in the methods section and Supplemental Information, Section 3. Due to the ambiguous nature of *C*_0_ in the context of electric fields from photons, particularly in relation to its ultrafast behavior, we opt instead to use this method to determine the shape result instead of any given magnitude in phase shift. The results of this model, some of which are shown in Figure 3, highlight differences between varying laser powers and beam waists.

**Figure 3.**
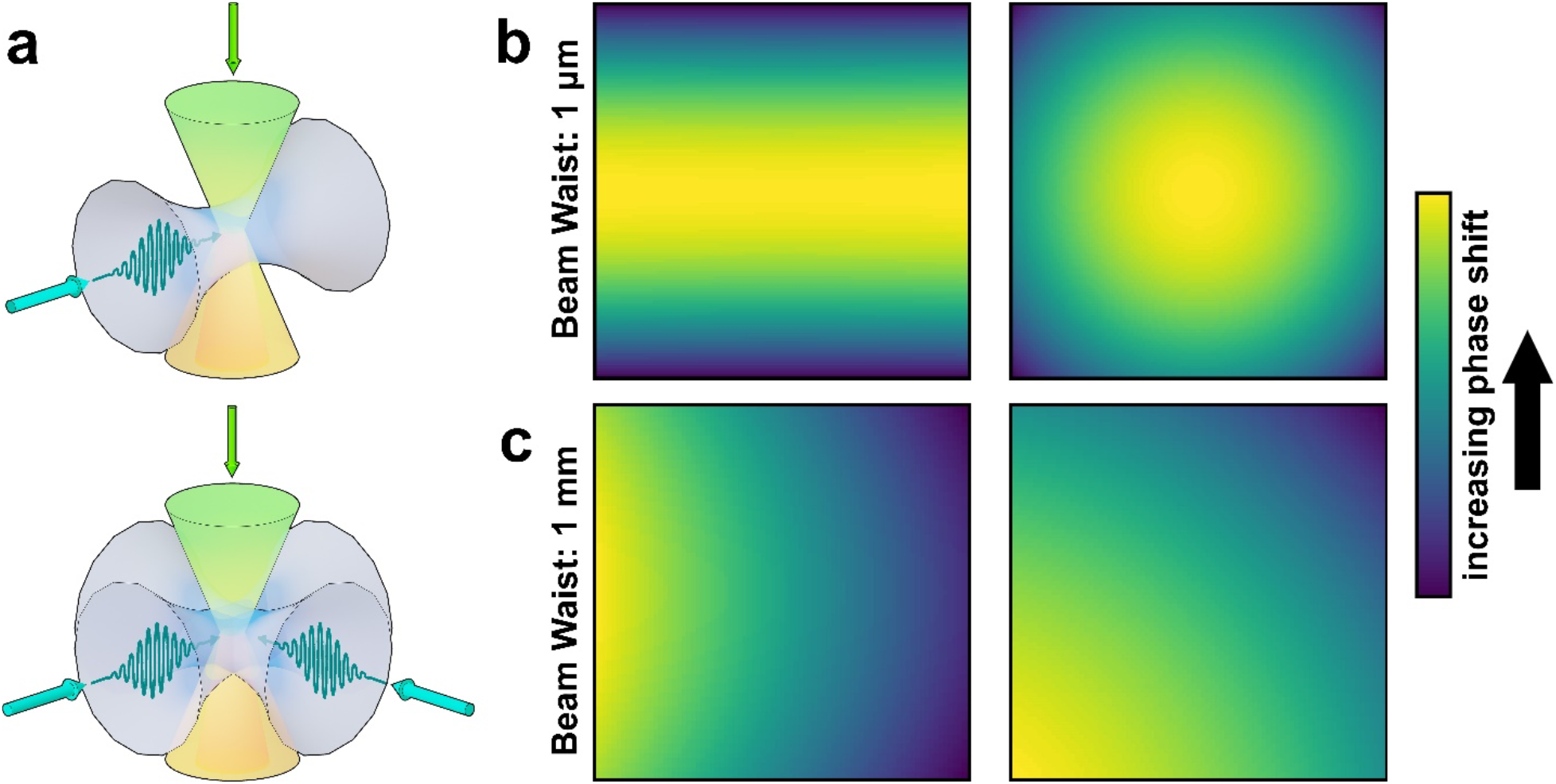
The contour plots displayed here are representative plots chosen from a wider range of tested experimental parameters and are meant to demonstrate the range of contours that appear from solving the quasi-classical model. Major variances in phase shift shape are a result of changes in beam waist size and no other simulated experimental parameters showed significant variations in the phase plate shape. A colorbar has been included at the right to show darker regions of lower phase shift and brighter regions with higher phase shift. (a) Schematic of one laser and two laser geometries in the ponderomotive phase plate with associated typical contour plots in the quasi-classical simulations. (b) corresponds to profiles with a laser beam waist size of 1 μm. (c) corresponds to solutions at laser beam waists of 1 mm.

The smearing effect expected from the interacting traveling waves is shown clearly in Figure 3. At smaller laser beam waists, the phase shift matches the results from the fundamental scattering model. However, by moving to the far more realistically feasible beam waist of 1 mm, we observe a stronger phase shift in electrons traveling in the same direction to the laser beam, an expected result for electrons that spend additional time in the high intensity regions of the laser beam. This smearing effect has been carefully checked by varying numerous simulation parameters and is further explored in the Supplemental Information. Nonetheless, we still do not observe any form of regular or repeating structure to the phase shift as expected from Kapitza-Dirac diffraction or as the mathematical form of the interaction equation the model suggests. Moreover, there is no change in the profile as the power increases, indicating that the smearing effects occurs regardless of the pulse energy.

One theory on mitigating the isotropic effects of a single laser has been to use two laser beams, positioned orthogonal to each other, such that the resulting phase shift forms rings or circles rather than lines (37). Here, we explore this hypothesis using an extension of both models by simply adding another laser source of equivalent characteristics oriented perpendicularly (*i.e*. traveling in the *x* axis) to the original source and with the same polarization and power, as shown in Figure 3a. The addition of a second laser only circularizes the phase shift in certain scenarios where the shift originally extends completely along the travel direction. However, Figure 3c shows in cases where the beam size is large and the resulting phase shift is prominently skewed towards electrons traveling alongside the laser travel direction, the expected circularization does not occur. More examples of this change in contour are presented in the Supplemental Information, Section 4. In no cases does there seem to be a reappearance in Kapitza-Dirac diffraction. From these simulations, we can preliminarily conclude that such a grating only appears in situations where the grating is static.

So far, these results have been discussed in the context of an electron beam that condenses to a singular or infinitesimally small point at the center of the laser beam (*i.e*. time-zero). It is important to note that while this provides a basis for which to continue discussions on the feasibility of a pulsed ponderomotive phase plate, the amplitude of any such phase shifts will be artificially inflated, as all electrons pass through the most intense region of the laser beam. To address this, another layer of the model must be added to include the effects of a finite-sized electron beam crossover.

### Size Effects in Both Approximations

Of course, an infinitesimally small crossover for the electron beam is an unreasonable expectation. In fact, in most electron microscopes, the direct beams are usually fit directly to a Gaussian or a Lorentzian as part of the instrument response function convoluted with the specimen response (38). We can take such a principle and apply it to the models provided above, where we perform simulations on pulsed beams passing through a region 200 μm across (a commonly available aperture size) and split into 30 sections, an arbitrary voxel size dictated by computational necessity, in both the *x* and *z* directions. Using a Gaussian with a similar width fit to 200 μm, each section is given a weight which is then used to calculate the average phase shift.

Figure 4 describes this addition to the calculation in more detail. Arrayed in the crossover region are the numerous simulation points denoted by cones that converge to infinitesimally small points. To join these simulation points together, a weighted sum is taken of each congruent angular direction (*i.e*. each independent travel direction), which either combine in the far limit or are reassembled at the image by additional lenses. For the sake of computational efficiency, the number of simulation points is diminished from the native granularity of each simulation. As such, each final point must include multiple “voxels” from the simulation matrices, as demonstrated in Figure 4a. The algorithm for this simulation is further explored in the Supplemental Information, Section 4. Here, we simulate the effect of a 1 nJ laser pulse interacting with an electron pulse during the span of a single nanosecond. The interaction region for the laser is 100 μm in beam diameter at its waist.

**Figure 4.**
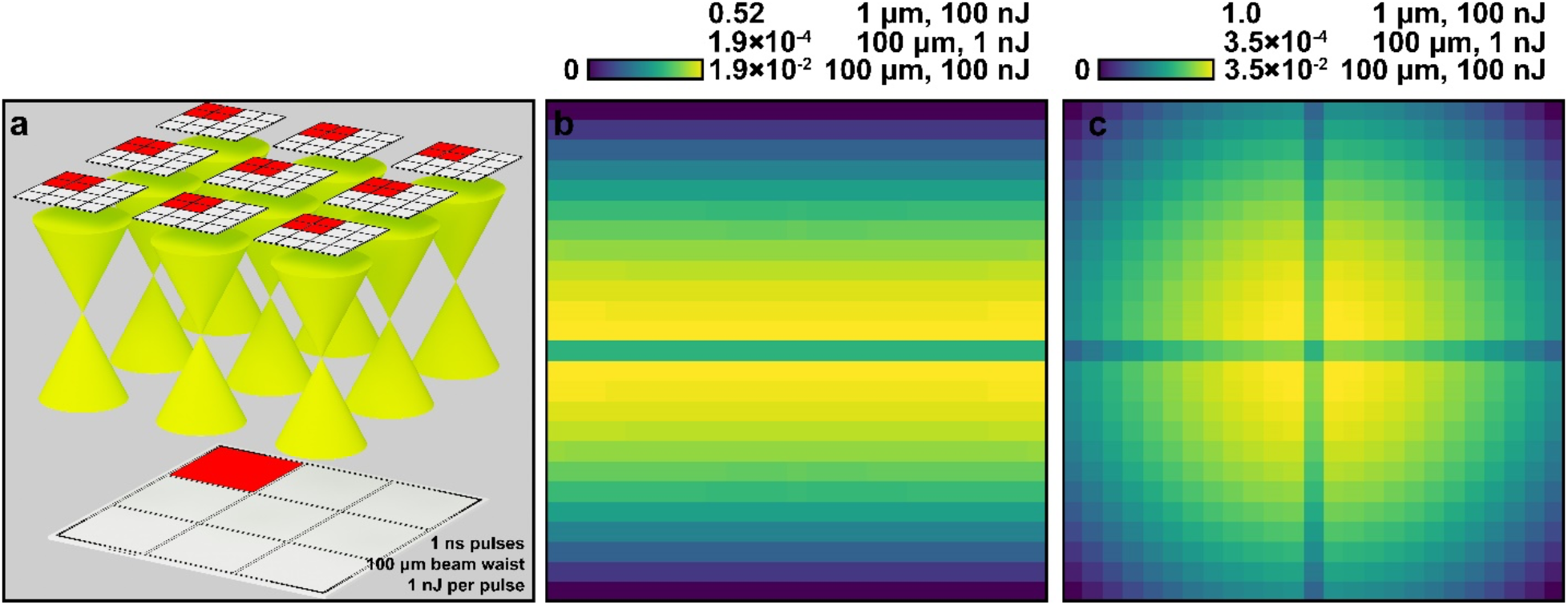
Demonstration of accounting for the finite size of the direct electron beam. (a) shows the simulation technique used to account for this non-infinitesimally small crossover at specific laser parameters. (b-c) correspond to phase shifts using the fundamental scattering model using a single and double laser respectively. (d-e) use the quasiclassical model. (f-g) show the combination of the scattering model and the normalized quasiclassical model, with colorbars (units in radians) to quantify the resulting phase shift.

By including the effects of size, we immediately observe a decrease in the expected phase shifts. This is an expected result, as the effects of the ponderomotive force are spread out into a finite region rather than a singular point for each electron to pass through. Interestingly, both the fundamental scattering and quasiclassical models converge to the same shapes, regardless of the underlying interaction equations. Three separate beam waist and pulse energy conditions are tested: 100 nJ at 1 μm, 1 nJ at 100 μm, and 100 nJ at 100 μm. The resulting phase shifts for a single laser beam are, respectively, 0.50 radians, 1.8 × 10^-4^ radians, and 1.8 × 10^-2^ radians. This suggests a linear relationship to pulse energy, though more calculations are required before completing this assessment. By taking the more realistic situation of using a 100 μm standard deviation beam waist laser, the required π/2 radians is reached at 8.7 μJ of pulse energy. Importantly, the peak instantaneous fluence attained by this pulse is approximately 2.2 GW/cm^2^, similar in fluence to continuous beam ponderomotive phase plates (30). This is attainable by Class 4 pulsed lasers and, at repetition rates of 100 kHz, we hypothesize 870 mW of power is required to attain the desired phase plate. Higher repetition rates may be required to reach the rapid acquisition rates of modern cryo-electron microscopy techniques, though such increases in repetition rate may lead to deleterious effects on the specimen (39). Nevertheless, the powers required from these simulations closely match in magnitude to the required input powers by other laser-driven phase plates (30). Indeed, we demonstrate a similar principle by temporally compressing the laser beam while other phase plates rely on resonance cavities to attain high Q factors in the electron optical axis (21, 31).

It must be noted that while the results of the quasiclassical model match those of the fundamental scattering model as shown in Figure 5, the simulation itself is greatly restricted by the number of points and mesh size in the *xz*-direction. Ultimately, at a crossover size of 200 μm, there should be at minimum 400 points to match the wavelength of the photons used in the simulations. However, computational limits greatly hinder this type of calculation and, in this case, only 25 calculation points along each axis were used. Ultimately, using a diminished number of points leads to aliasing and a shape that is not truly representative of the solution, as demonstrated by the anomalous dip in phase shift at the center of each contour plot. Nonetheless, a brief study in increasing the mesh sizes demonstrates that while the number of points is unsatisfactory, the resulting shapes are sufficiently close enough to the non-aliased solution. Additional information on this is provided in the Supplemental Information Section 6.

**Figure 5.**
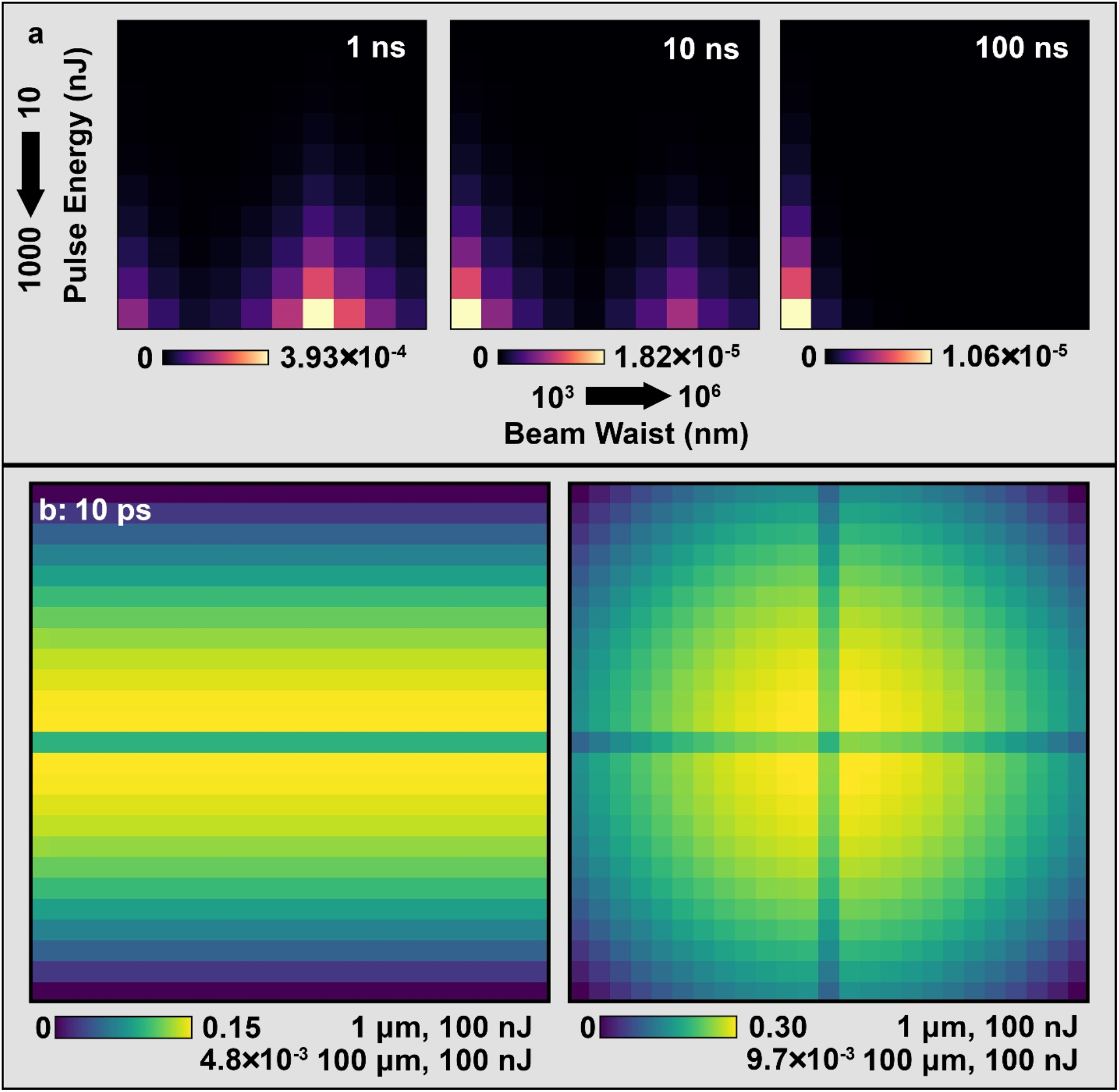
An exploration of the interaction between a uniform (*i.e*. stochastically emitted) electron beam and pulsed laser beams. (a) demonstrates the peak phase shift as a function of pulse energy, beam waist, and pulse duration of the laser. These simulations used the infinitesimally small approximation. (b) shows the simulation results in the presence of a finitely sized electron beam, with the peak phase shifts from two cases displayed next to each colorbar. The single laser case is on the left and the double laser case is on the right. The fundamental scattering and quasiclassical models converged to the same phase contour at these simulation resolutions.

### Stochastic/Uniform Electron Emission

Before the implementation of pulsed electron beams, it is important to verify whether a strong enough ponderomotive effect is possible in a continuous electron beam. While pulsed beams are compelling for the level of control in emission timing, the deleterious effects from bunching charges is well known (40). Even in the single-electron regime, variations in hardware at the cathode can greatly affect the emission behavior of electrons and introduce aberrations (41). As a result, the most ideal condition for the ponderomotive force to occur would be in the native unmodified electron beam. Of course, while the emission of electrons is largely stochastic in nature when not under ultrafast conditions, the beam can be considered as continuous over longer acquisition times (39, 42–44).

We simulate such a beam by flattening the probability weights of each slice such that any electron has an equal probability of appearing in each slice. The results of this uniform electron emission, presented in Figure 6, can then be appropriately adjusted to account for the percentage of time that there is no ponderomotive force in the column:

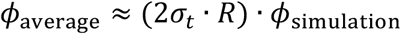

where *R* is the repetition rate of the laser and *ϕ*_simulation_ is the phase shift calculated in the duration of the ponderomotive force. Ultimately, the necessity of containing a significant fraction of time largely limits most laser capabilities to the nanosecond range, as there are few commercially sold pulsed lasers that operate on the femto- or pico-second timescales while still maintaining the necessary repetition rates for efficient temporal coverage of a uniform electron beam (*i.e*. 100 GHz for 10% coverage in a picosecond laser and 1 THz for 50% coverage in a 500 femtosecond laser). We limit our analysis to nanosecond lasers operating in the much more reasonable MHz range.

**Figure 6.**
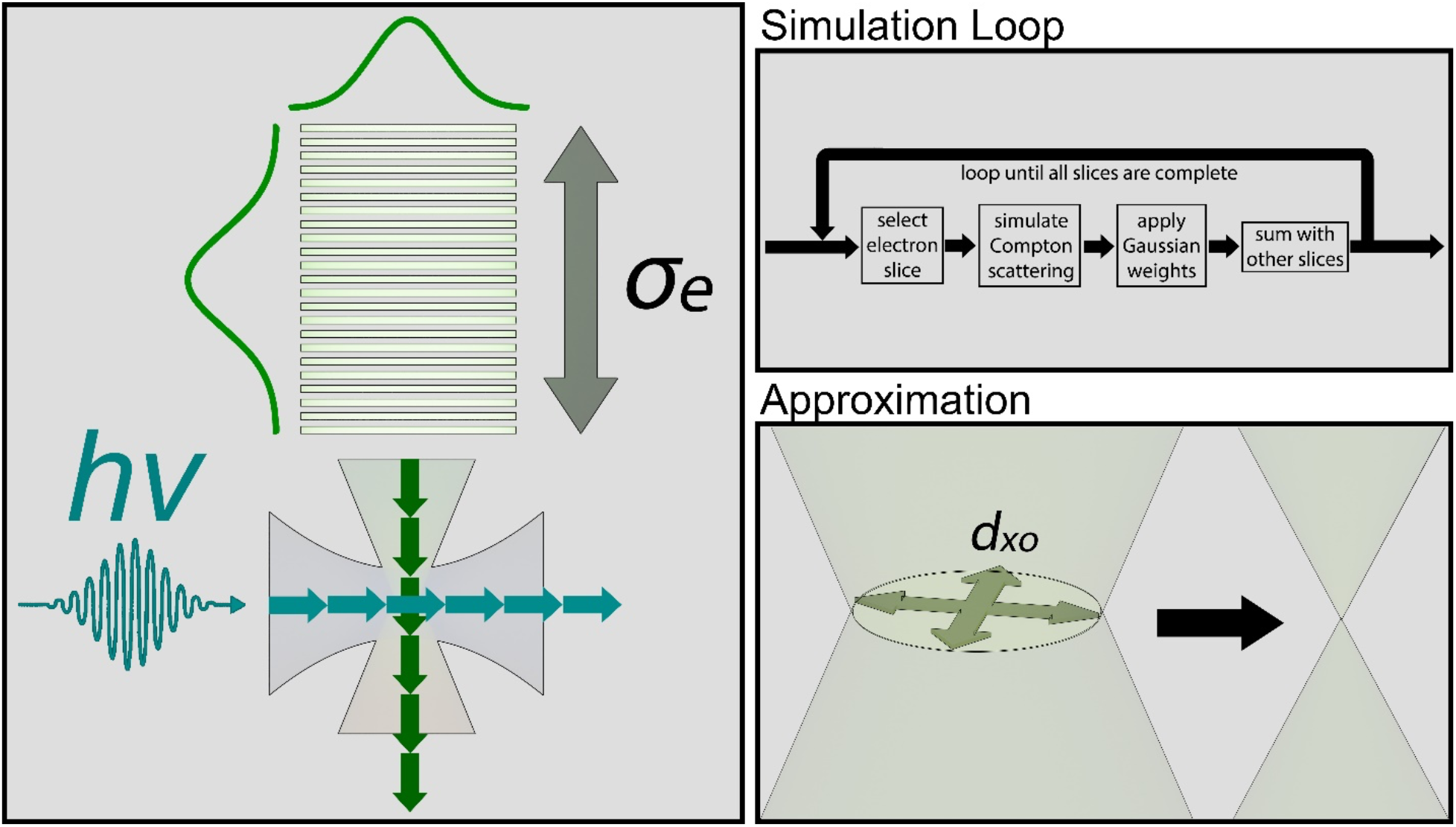
A schematic of the forward-stepping finite element model algorithm. The electron beam is split into slices of varying thickness and stepped through the interaction region. The full algorithm is displayed on the top right, showing that each of the electron slices has a Gaussian weight in both the spatial and temporal directions. These weights are applied at the end of each interaction region and summed with the results of every other electron slice to determine the average phase shift in the field of view. A key assumption made in all but the last section of this paper shows the electron crossover being reduced to zero, forcing all electrons to pass through the densest region of the laser beam.

Figure 5a clearly shows the peak phase shifts in the uniform electron beam case before adjustments to the proportion of time the laser is in conta
ct with the electron beam. There is no experimental situation where the electron beam experiences a phase shift by π/2 between 1 and 100 ns of laser pulse duration. When accounting for the case of a finite electron beam focal point, the phase shifts become even less promising, with peak phase shifts of 0.30 radians at dual beam waist of 1 μm, powers of 100 nJ, and pulse durations of 10 ps. The pulse energies required to attain π/2 phase contrast rapidly exceeds hundreds of μJ per pulse. At MHz frequencies, such powers become unfeasible.

## Methods

As described previously, these models were adapted from previously published literature describing the phase shift in Compton scattering and fit to a forward-stepping algorithm to determine the influence of ultrafast pulses on the phenomenon (21, 30). More specifically, the electron beam is split in to slices and each slice is individually stepped through time as it passes through the laser beam, beginning three laser pulse spatiotemporal radii (3σ, 99.7%) away from the center of the intersection region. The crossover slope for the electrons is set to be over the span of a foot (300 mm). At the end of each slice calculation, it is averaged into the total amount of slices that form the full electron pulse, as described in Figure 7. Numerous concessions were made to improve simulation times, including ignoring the electron beam diameter at the crossover point (*d_xo_*) and a loss in granularity of the forward-stepping algorithm (121 side-length voxel cube). It is important to note that this pinched crossover artificially increases the phase shift of the electrons and is the cause for the section dedicated to size effects in both models. Finally, we assume a noninteraction regime between electrons. Additional information on this is provided in the Supplemental Information Sections 1, 2, and 5.

## Acknowledgments

This material is based on work supported by the Chan-Zuckerberg Initiative, Visual Proteomics Grant under award No. 2021-234816.

## Supporting Information

### 1. Ultrafast Gaussian Pulse

In the main text, we described the behavior of an ultrafast gaussian pulse as following the equations (1):

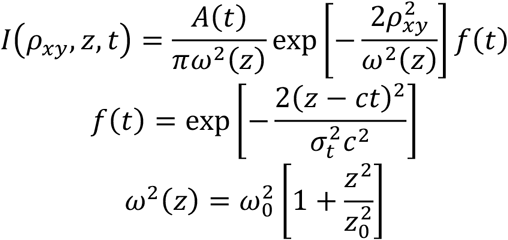

where *A*(*t*) is a time-varying parameter used to normalize the pulse to the pulse energy, *ω*(*z*) is the gaussian beam diameter, *z*_0_ is the Rayleigh range, *ρ_xy_* is the radial distance from the travel axis of the laser, *σ_t_* is the time resolution, *c* is the speed of light, and *z* is the distance down the travel axis.

**Figure S1.**
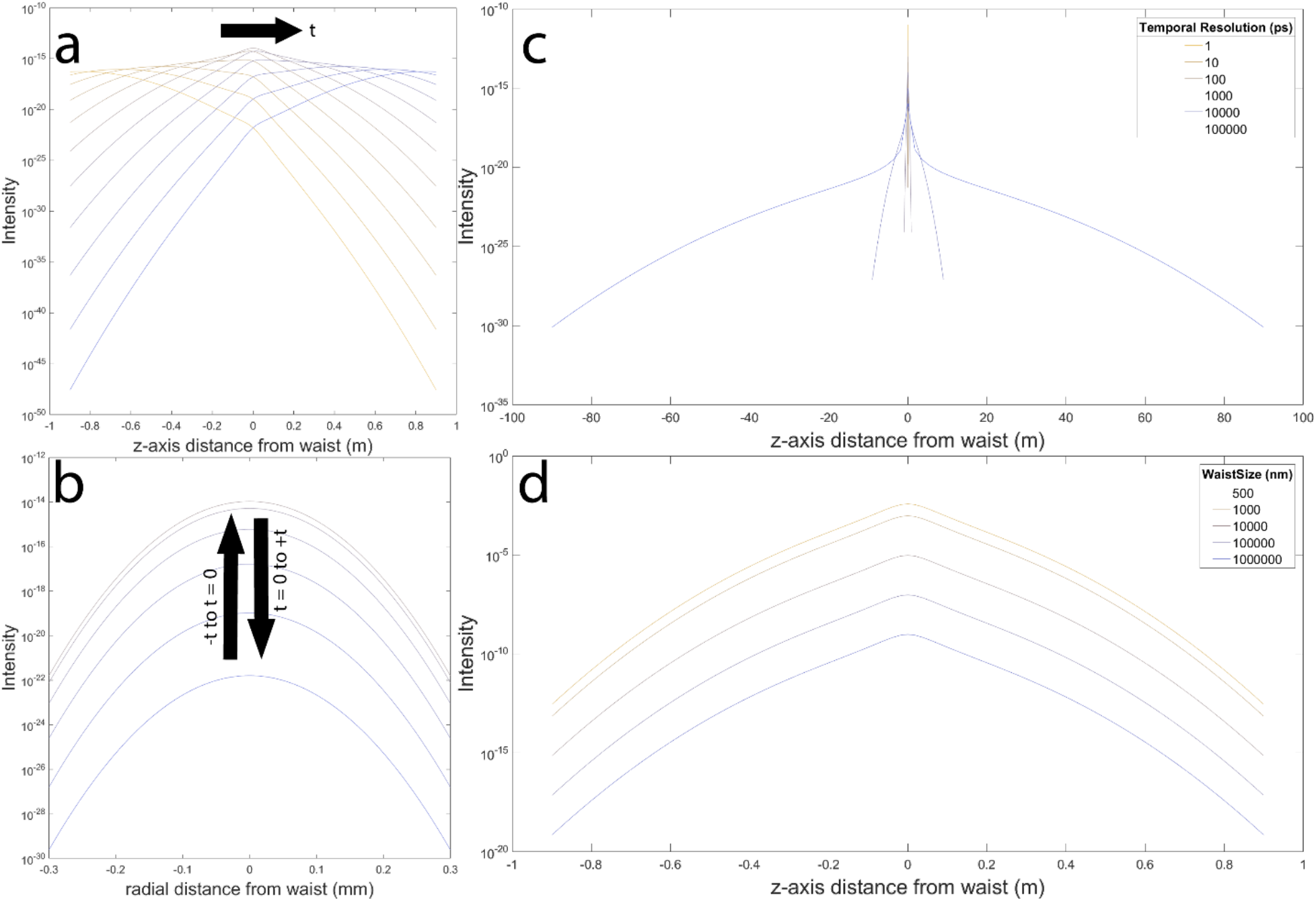
A demonstration of the behavior from the ultrafast pulsed laser equation. (a) shows a beam with a beam waist of 100 μm and a temporal resolution of 1 ns. The beam travels from left to right (yellow to blue). (b) shows the same beam in the radial direction at *z* = 0 along the propagation axis. The intensity initially increases before decreasing in a temporally symmetrical manner. (c) A comparison of temporal resolutions ranging from 1 ps to 100 ns with a beam waist of 100 μm. Sharper temporal resolutions give rise to sharper increases in intensity along the *z*-axis. (d) A plot of different beam waist sizes ranging from 500 nm to 1 mm with a temporal resolution of 1 ns. The smaller the beam waist, the higher the intensity along the *z*-axis.

It is important to note that a true simulation of an ultrafast pulse involves a traveling wave of multiple wavelengths in a certain dispersion around a central wavelength. Here, we simulate this pulse by applying a temporal component in two places of the original Gaussian beam equation. This is an approximate solution for simulation purposes and is not an analytical solution to the ultrafast laser pulse.

### 2. Full Simulation Algorithm

This paper involved the use of 16 separate scripts. These scripts can loosely be grouped into two categories: those that account for the size of the crossover and those that do not. In this section, we will discuss the algorithm that is used to perform the calculations for the zero-crossover case. This algorithm ultimately forms the basis for the nonzero-crossover case.

**Figure S2.**
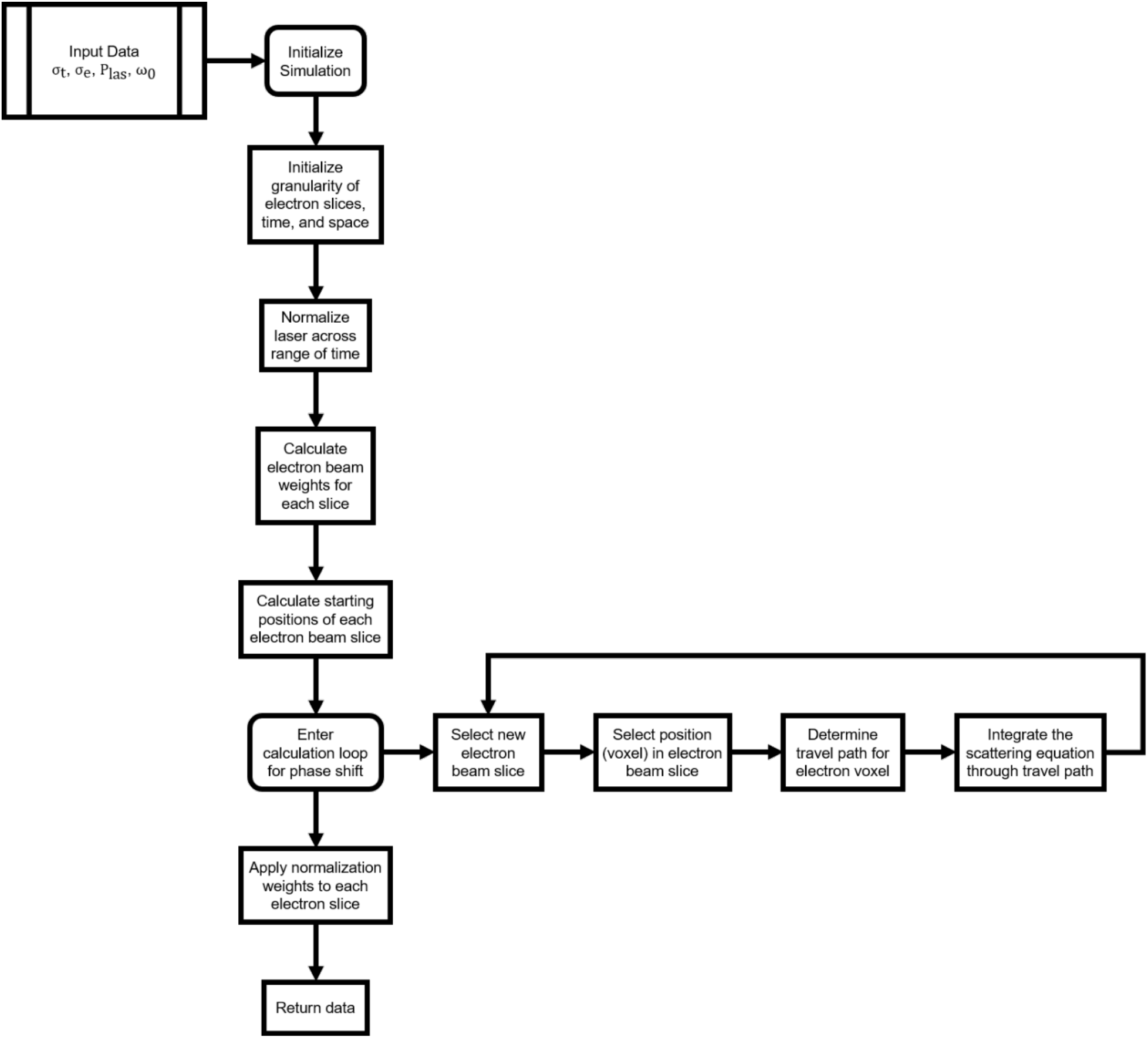
An algorithm diagram providing an in-depth description of how the simulation for the zero-crossover cases are handled.

### 3. Fundamental Scattering Model

Here, we explore the electron slices in the scattering model as a demonstration of how the phase is distributed. We have split each figure into two parts: one with the peak phase shift labeled at the top and one with the time that the electron beam reaches the center of the laser beam.

The first set involves 10 ps pulse duration beams while the second set involves 1 ns pulse durations.

10 ps pulse durations:

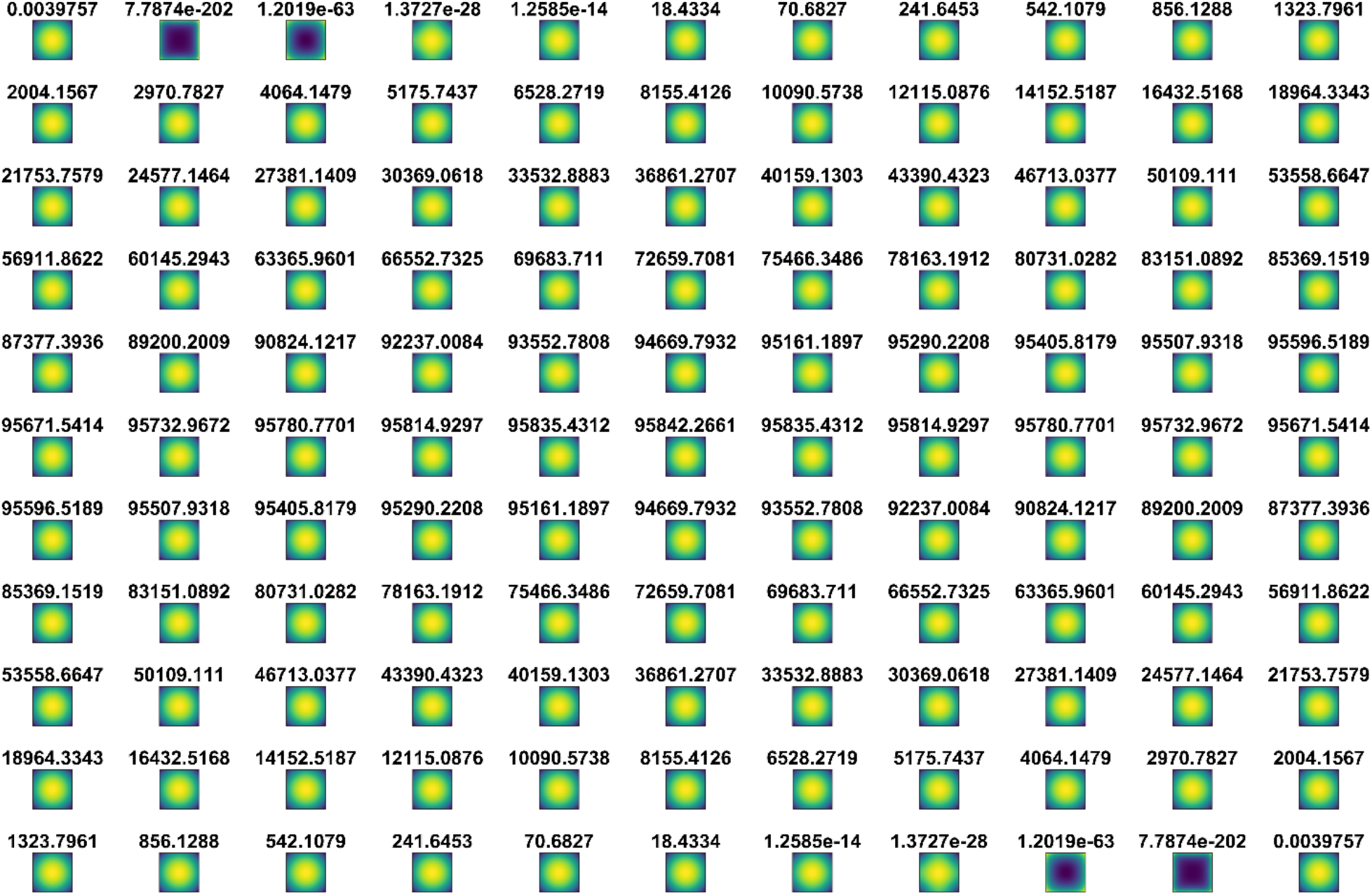
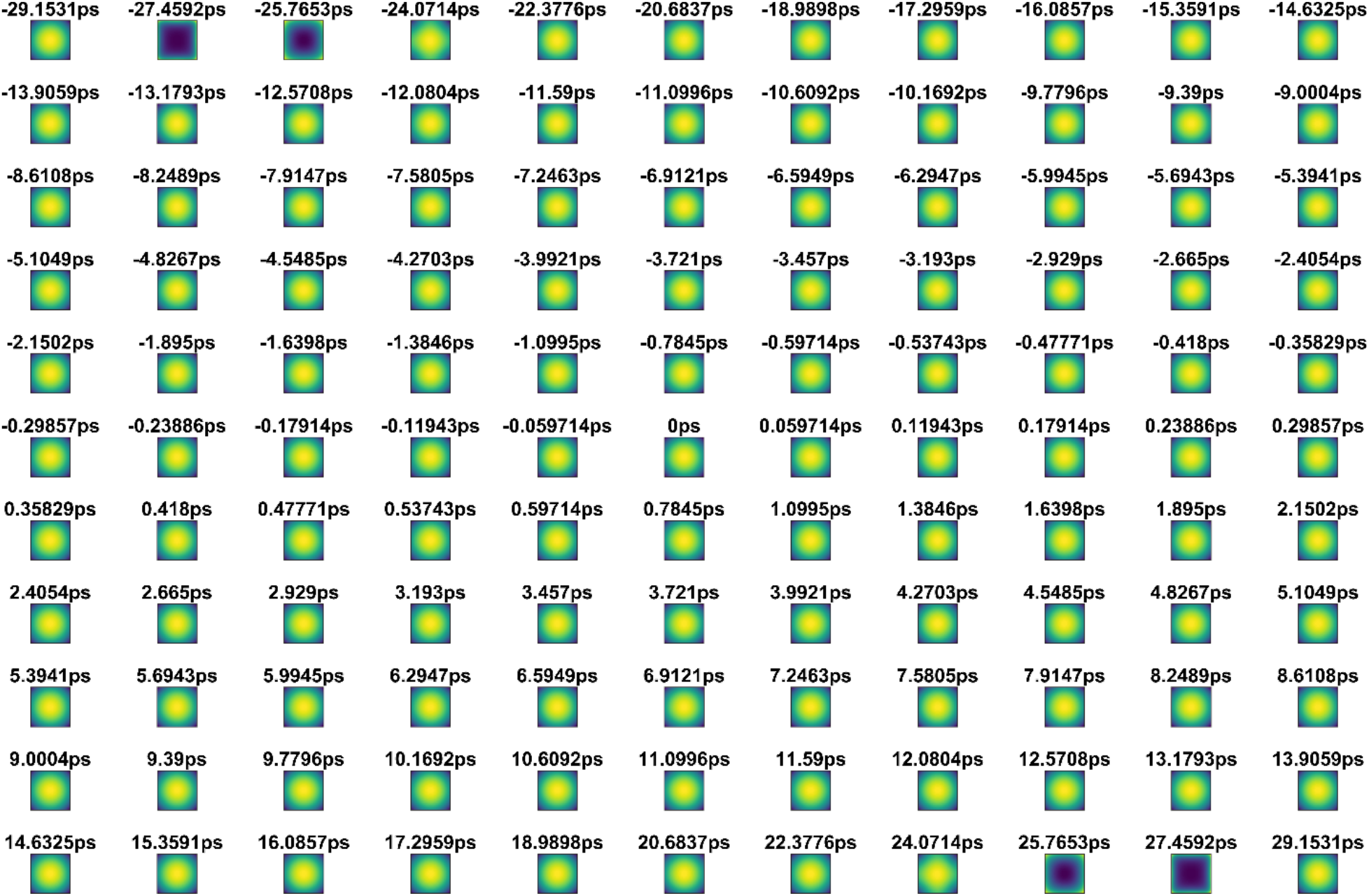

1 ns pulse durations:

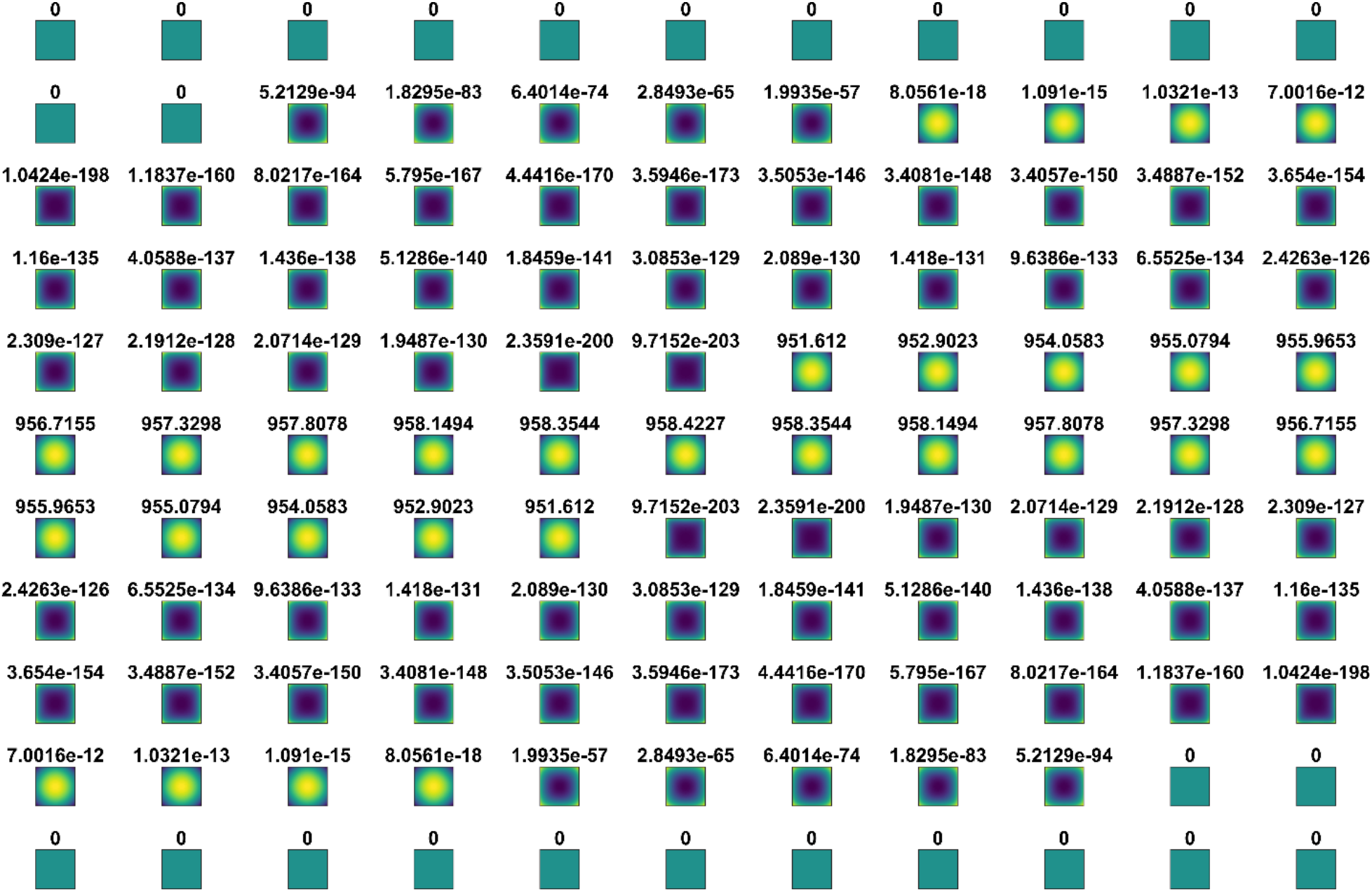
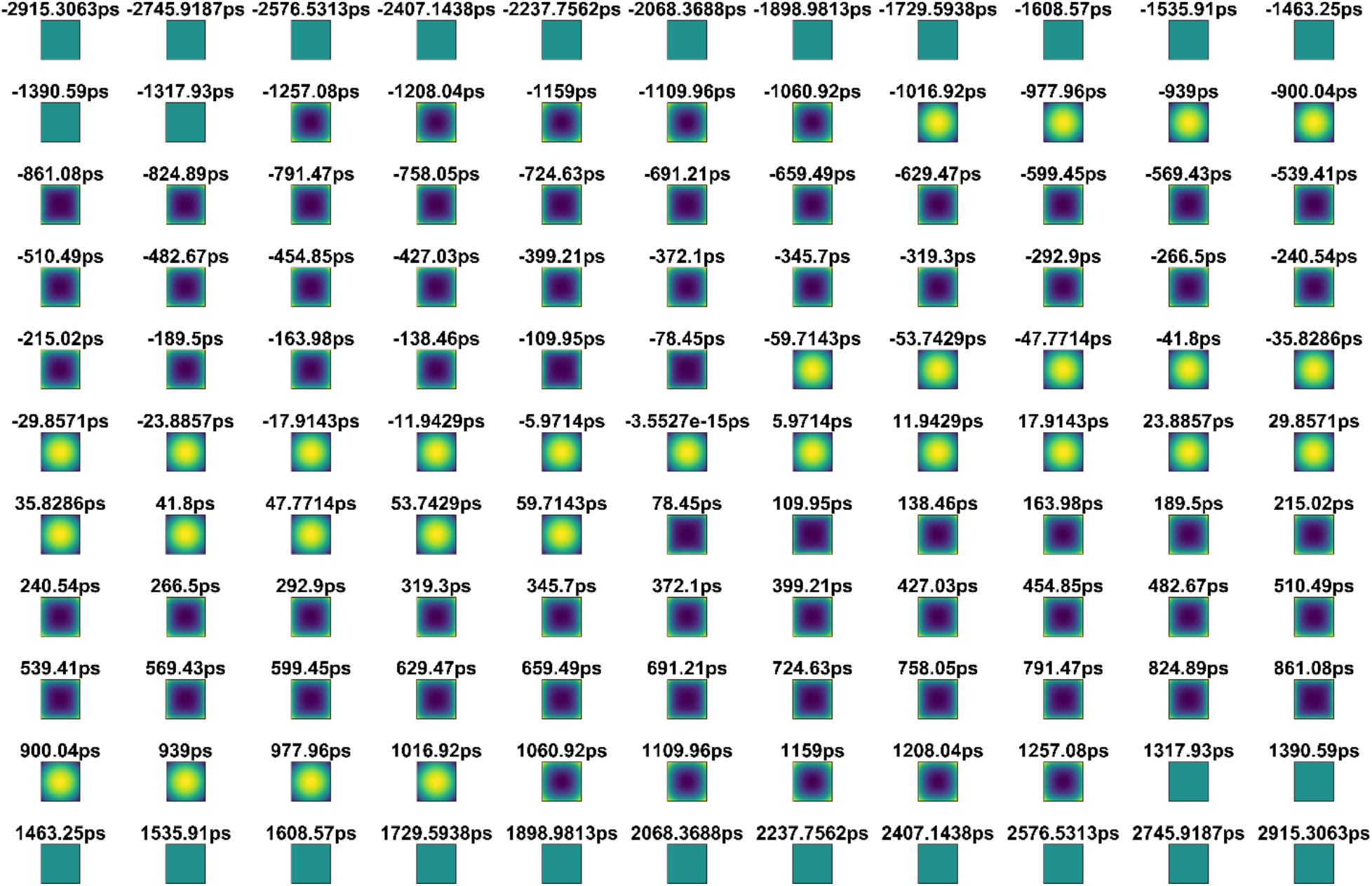

### 4. Quasiclassical Model

In the main text, we used the following equation as the foundation for the quasiclassical simulation model, based heavily upon work done by Turnbaugh *et. al*. (2):

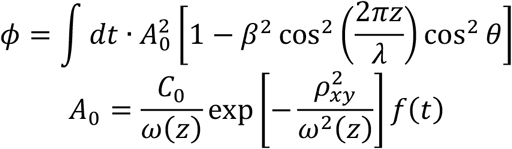

where *C*_0_ is an arbitrary vector amplitude, *β* = *v*/*c*, and *θ* is the polarization of the laser (here set to zero). Here, we will go through the derivation of the above equation beginning with the generalized gauge invariant solution to the phase shift as presented by Turnbaugh *et. al*.

Derivation:

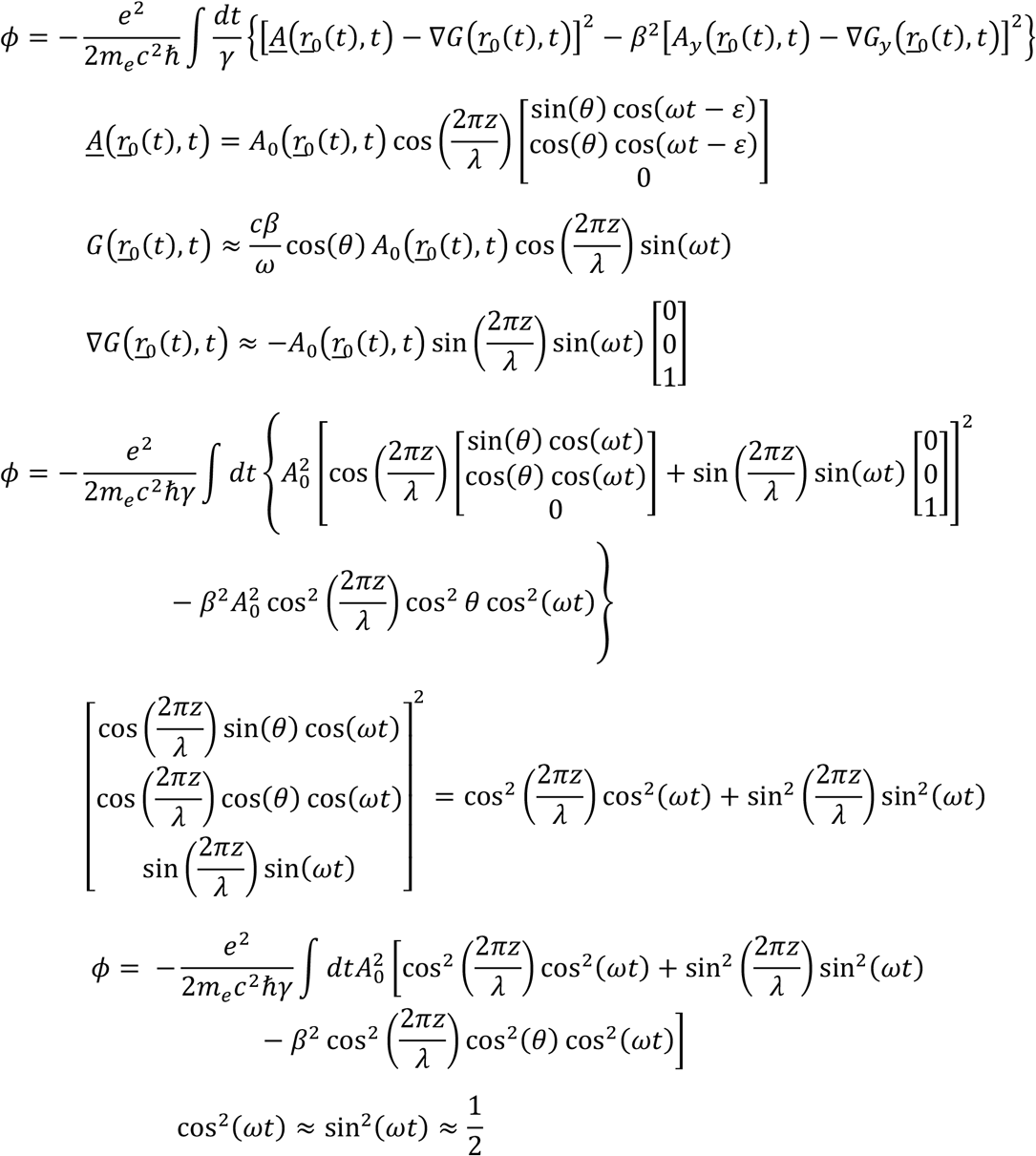

which is taken as a 0^th^-order average of the two trigonometric functions due to the rapid oscillatory behavior from *ω*:

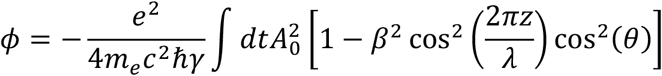

All that must be done now is to normalize the laser pulse at every point in time to some constant value. Given the units of *C_0_* are in Volts in a calculation involving photons, this leaves no recourse but to use arbitrary values as the normalization constant, labeled in the main text as relative power.

#### Examples of Results

The dual-laser solutions to the model above have been included here. They are segmented into individual electron slices without an applied normalization weight. At 10 ps pulse durations, the only major variations occur at different beam waists.

1 micron beam waist:

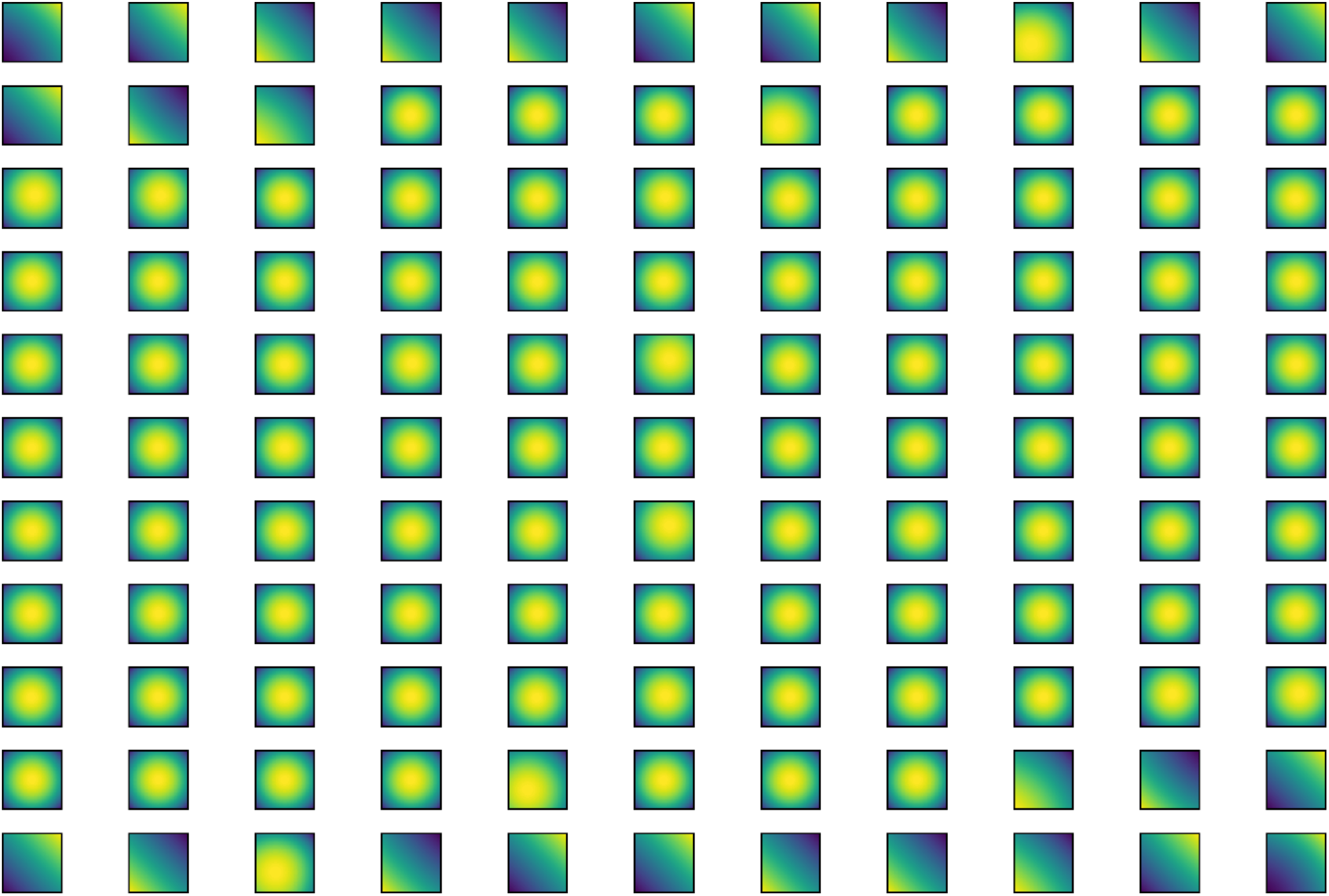

100 micron beam waist:

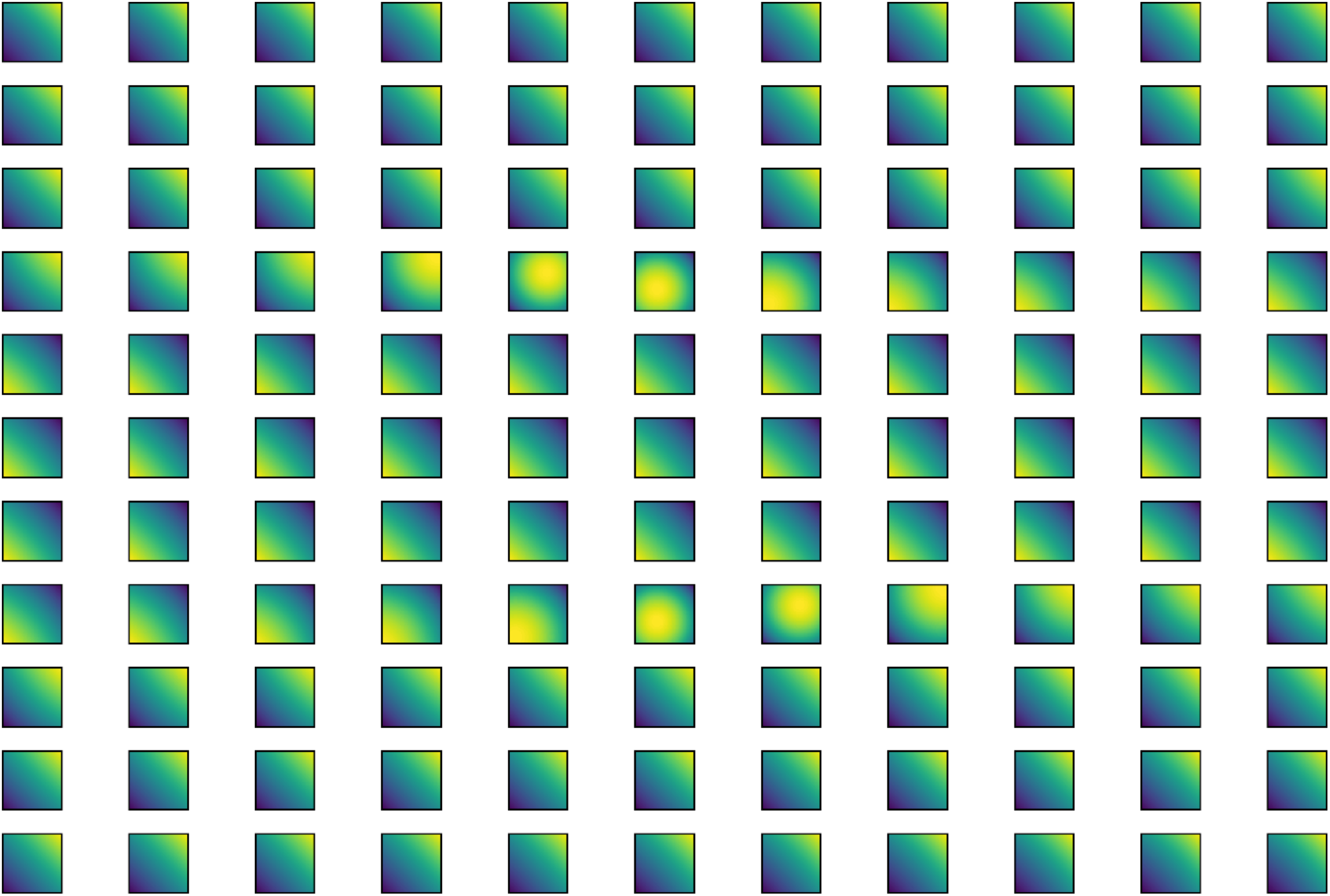

### 5. Simulation Algorithm to a Non-Infinitesimally Small Crossover

The algorithm for the nonzero-crossover case includes a spatial loop that covers the original temporal electron slice loop from the zero-crossover case.

**Figure S3.**
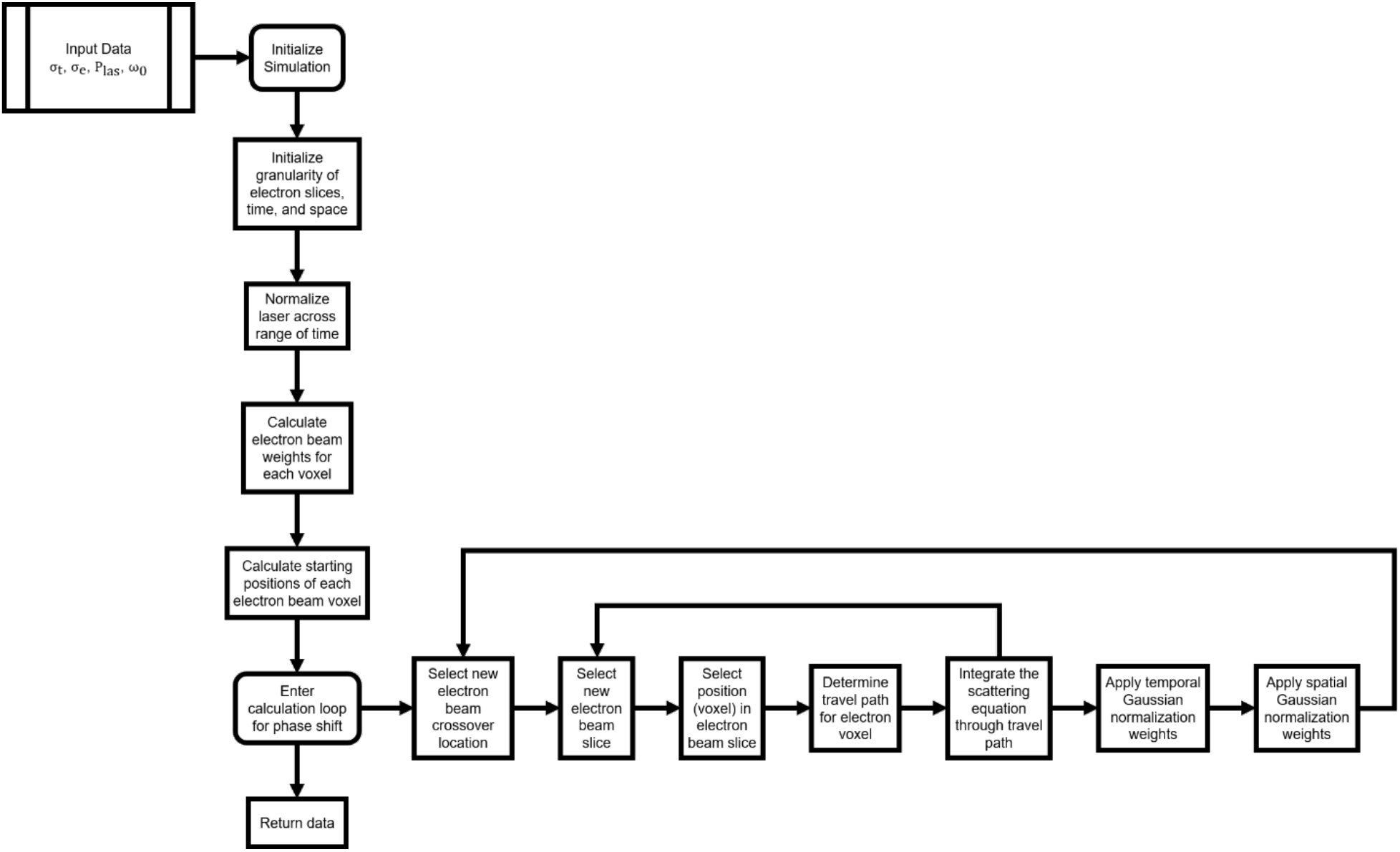
An algorithm diagram providing an in-depth description of how the simulation for the nonzero-crossover cases are handled. Here we see that a secondary loop has been included on top of the original zero-crossover loop to handle a shifting electron beam, thus simulating a real size to the electron beam crossover.

### 6. Distribution of Calculation Points and Mesh-sizing

#### Distribution of Calculation Points

The simplest form of distributing calculation points in the mesh of a finite element model (FEM) is an even spread across time and space, agnostic to the geometry of the specimen or form of excitation to the system. However, this can lead to results that are poorly representative of the true solution. To curb any skewing of the data, a weighted distribution of points is often generated that concentrates more calculation points towards regions of the system where interactions are more likely to occur.

In the context of the pulsed ponderomotive phase plate, the regions of interest would be when the photon pulses and electron beam meet and fully overlap. Within the simulation itself, we split the electron beam into the mesh and use a mathematical formulation of the laser beam to calculate the strength of the interactions. As such, the weighted mesh contains the density of electrons within the electron pulses. For the infinitesimally small crossover calculations, the mesh in the *xz*-plane, or the plane normal to the travel path, is distributed evenly as the incident electron beam is coherent and collimated from the illumination lenses. However, the distribution of points in the *y*-axis is weighted by the method in which the electrons are emitted. For an ultrafast photoexcited pulse, this shape would similarly take the form of a Gaussian.

For a finite crossover, the calculation becomes somewhat more complex. In the previous example, the distribution was assumed to be uniform due to the parallel illumination provided by the electron optics. However, in the case where we observe a finite-sized direct beam in the back-focal-plane, the distribution is closer to a Gaussian or a Lorentzian, depending on the instrument response function. Thus, in the final averaging of phases for electrons traveling at congruent angles, the final value must be weighted by the shape of the contour of the direct beam. In the paper, we mentioned this took the characteristic of a 200 μm aperture.

#### Mesh-sizing

The size of the mesh is also important. Obviously, a single data point cannot be used as the basis for the calculation. While the ideal number of points is infinity, such large array sizes are unreasonable and simply cannot exist with current memory capabilities. Thus, a series of tests were performed with weighted meshes to discover the ideal mesh size in the *x*, *y*, and *z* directions. The data is shown in Figures S4–S9.

**Figure S4.**
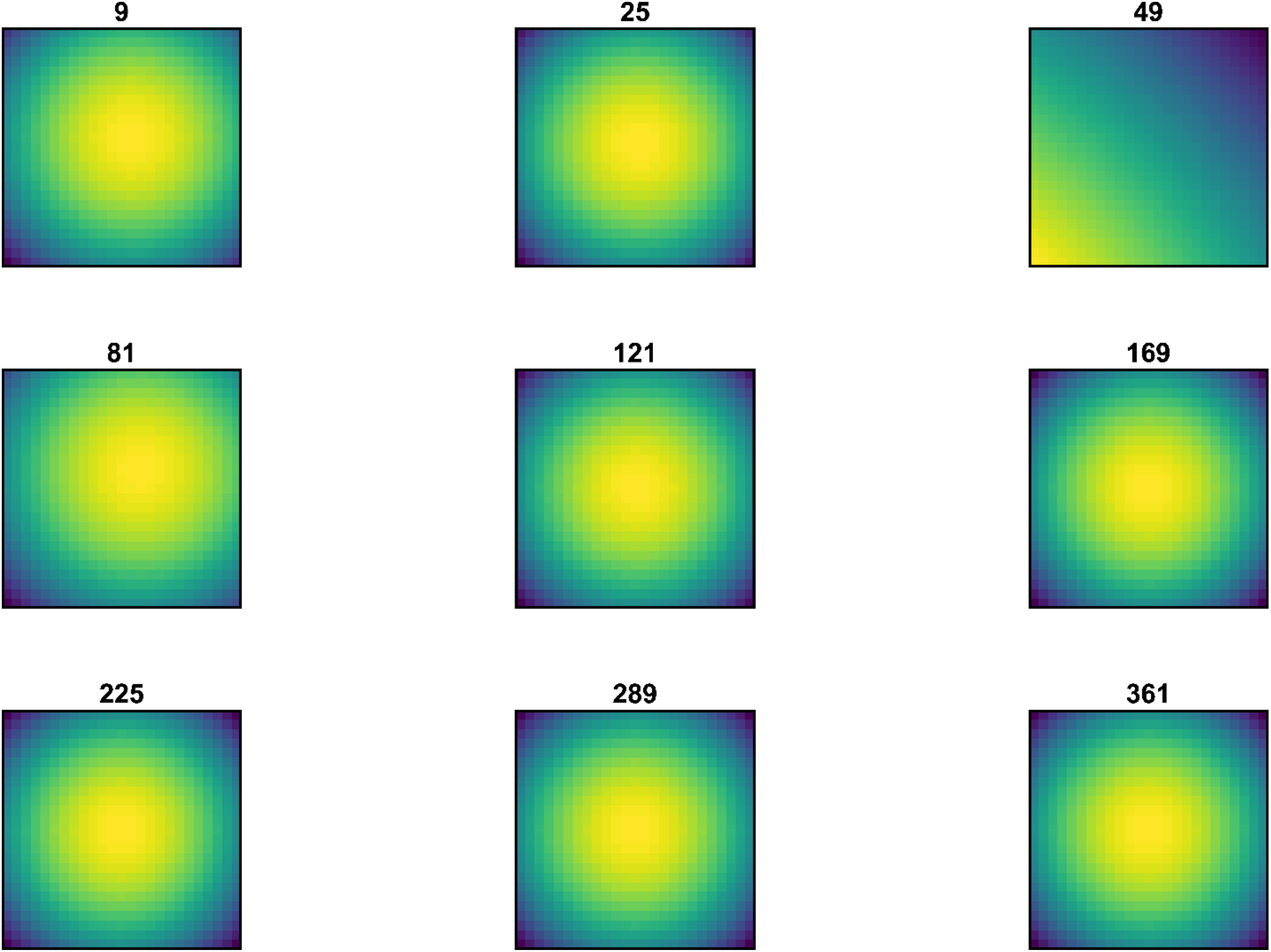
Quasi-classical model. Simulation conditions of 1 ps electron and laser pulses, 1 micron beam waist, and a relative power of 1000. The titles indicate the depth of the mesh size in the *y* direction. We clearly see an evolving phase shift profile as the mesh size increases and it does not appear to level off. Clearly more work is required to explore this phenomenon in depth, though the calculation times required to complete these simulations rapidly reach unreasonable requirements without the usage of dedicated supercomputing time. Only a single laser was used in this instance.

**Figure S5.**
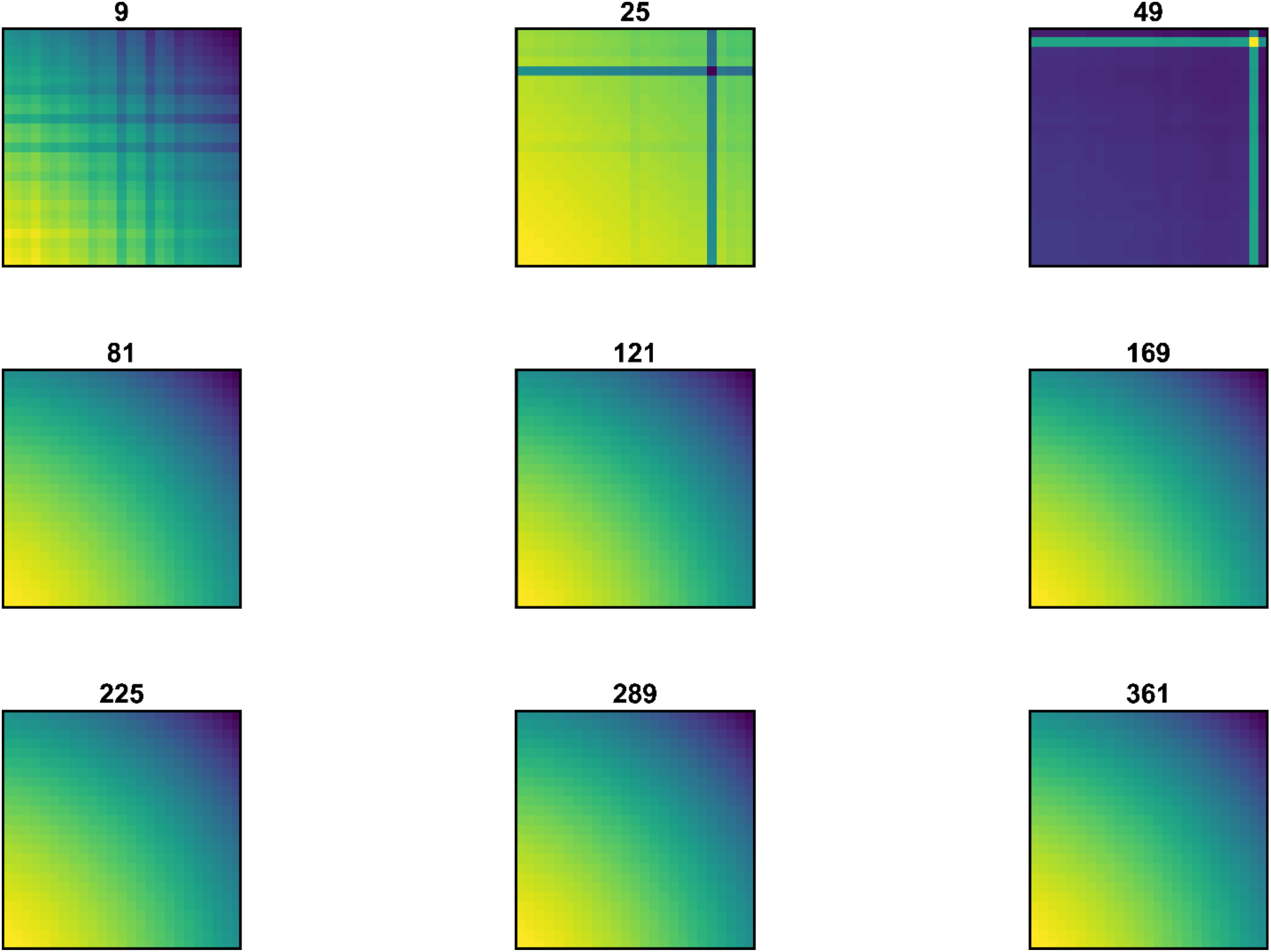
Quasi-classical model. Simulation conditions of 1 ps electron and laser pulses, 1 micron beam waist, and a relative power of 1000. The titles indicate the depth of the mesh size in the *y* direction. We clearly see an evolving phase shift profile as the mesh size increases and it does not appear to level off. Clearly more work is required to explore this phenomenon in depth, though the calculation times required to complete these simulations rapidly reach unreasonable requirements without the usage of dedicated supercomputing time. Double lasers were used in this instance.

**Figure S6.**
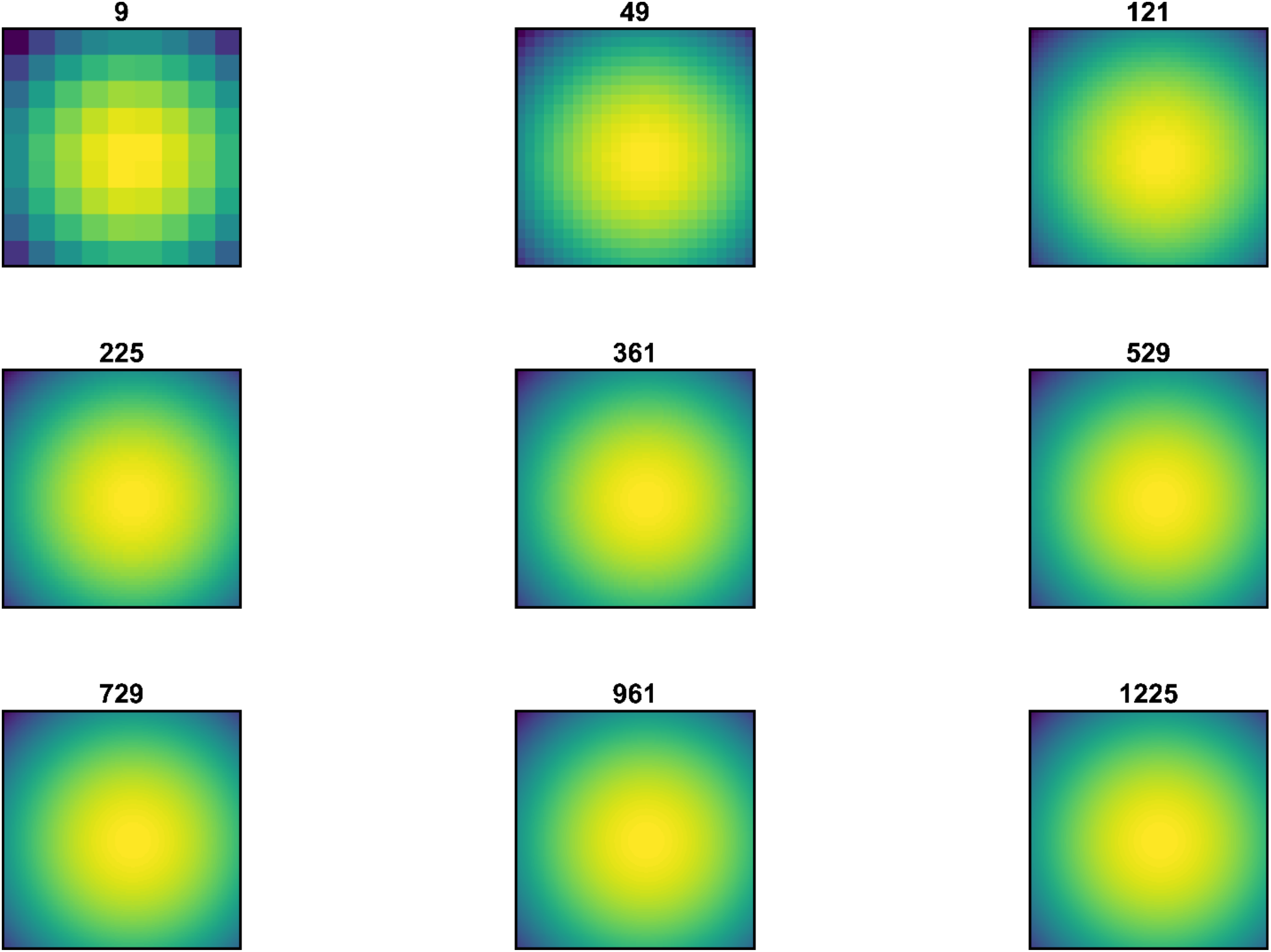
Quasi-classical model. Simulation conditions of 1 ps electron and laser pulses, 1 micron beam waist, and a relative power of 1000. Here, the effects of increasing the number of calculation points in the *xz* (indicated by the first coordinate) directions and the *y* (second coordinate) direction is demonstrated. We clearly see that when the *xz*-mesh is increased in size, there is a minimal effect on the shape of the phase shift. However, an increase in the mesh size in the *y*-direction shows a dramatic difference. Two lasers were used in this instance.

**Figure S7.**
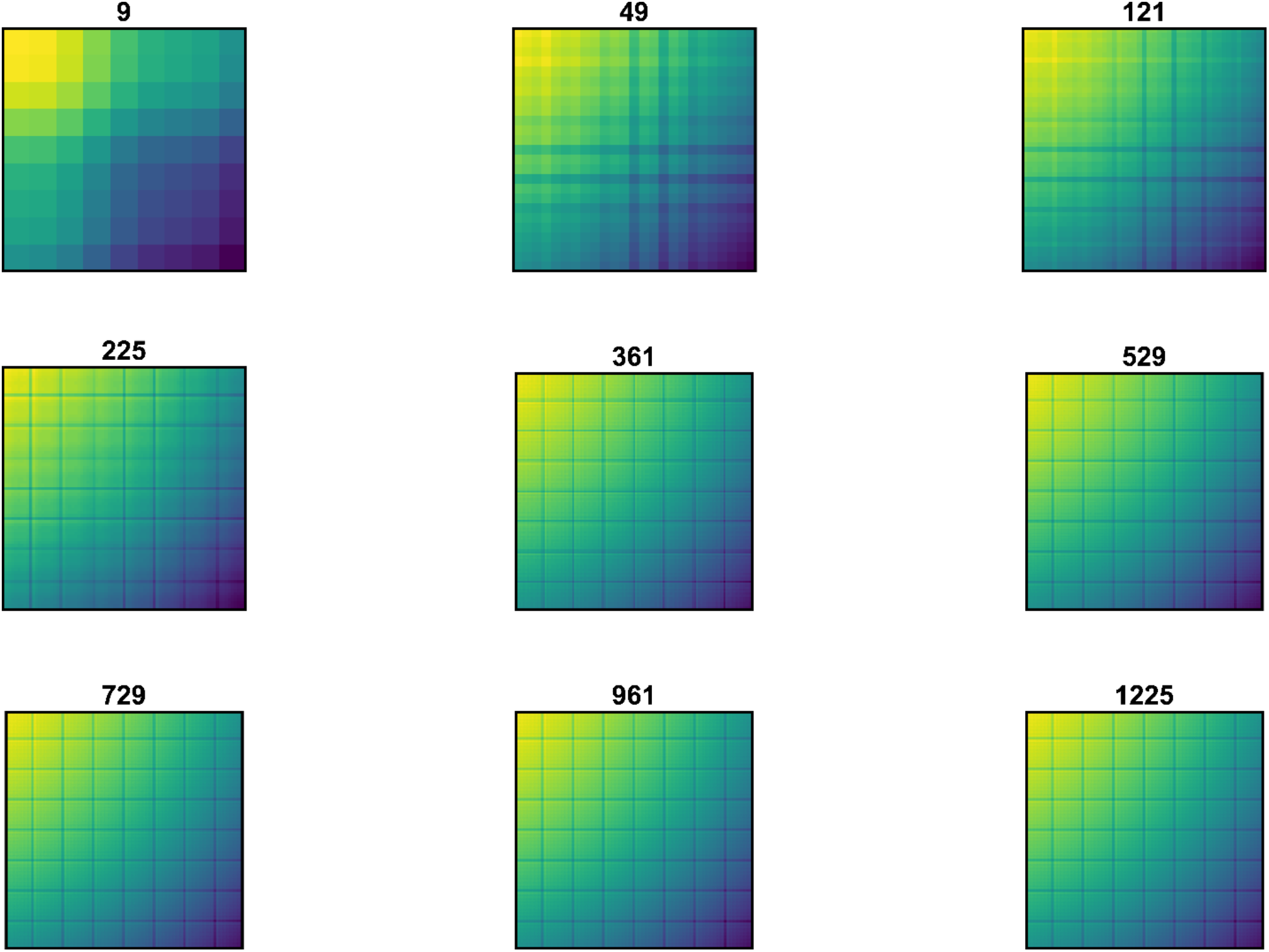
Quasi-classical model. Simulation conditions of 1 ps electron and laser pulses, 1 micron beam waist, and a relative power of 1000. Here, the effects of increasing the number of calculation points in the *xz* (indicated by the first coordinate) directions and the *y* (second coordinate) direction is demonstrated. We clearly see that when the *xz*-mesh is increased in size, there is a minimal effect on the shape of the phase shift. However, an increase in the mesh size in the *y*-direction shows a dramatic difference. Only a single laser was used in this instance.

Due to the non-oscillatory nature of the fundamental scattering model, the values converge much more quickly. As such, a smaller range of mesh sizes in all directions were tested and are presented here.

**Figure S8.**
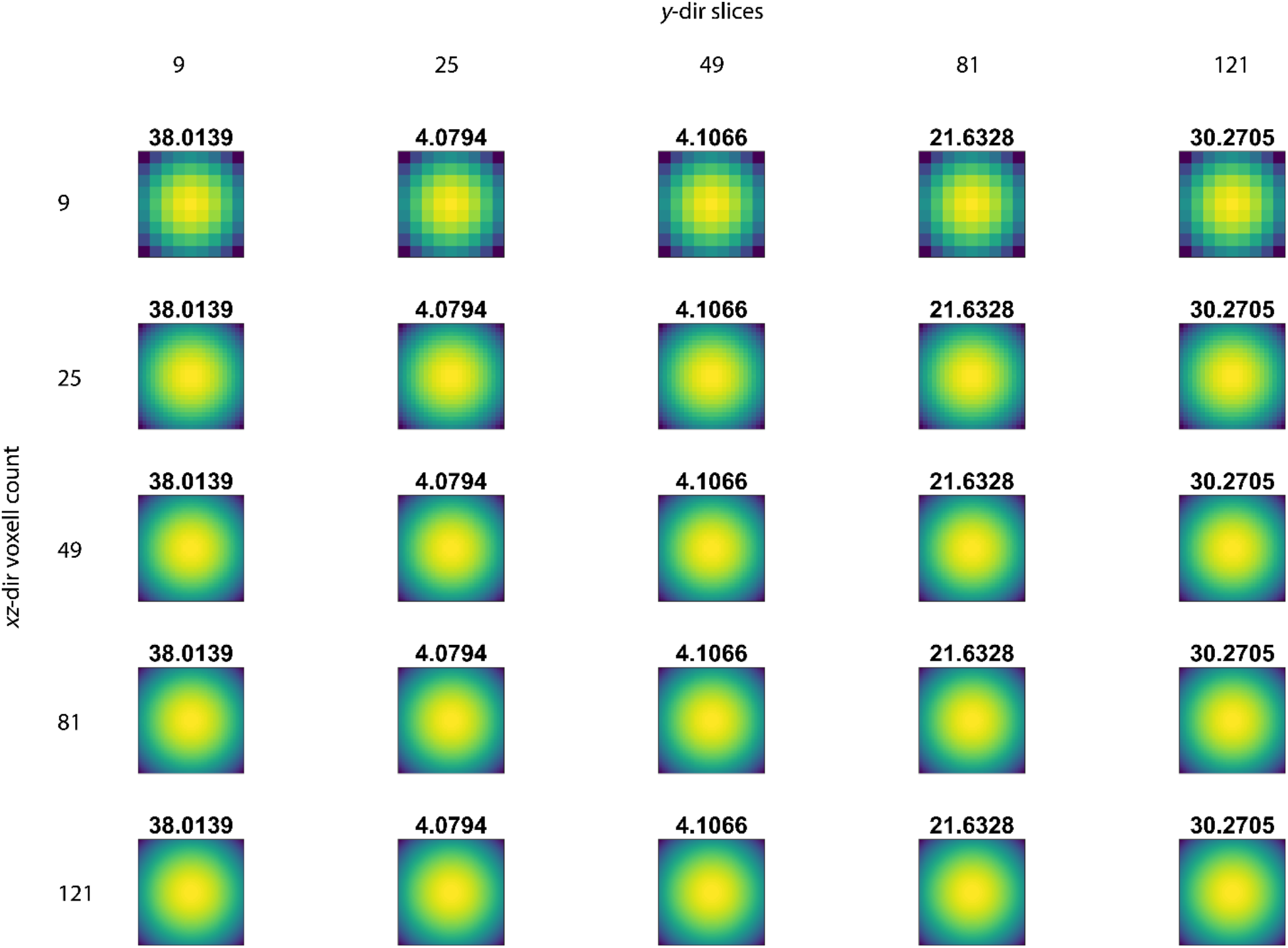
The values for the fundamental scattering model converge at around 121 *y*-slices to the reported values in the main text. Changing the voxel count in the *xz* plane does not affect the reported phase shift.

**Figure S9.**
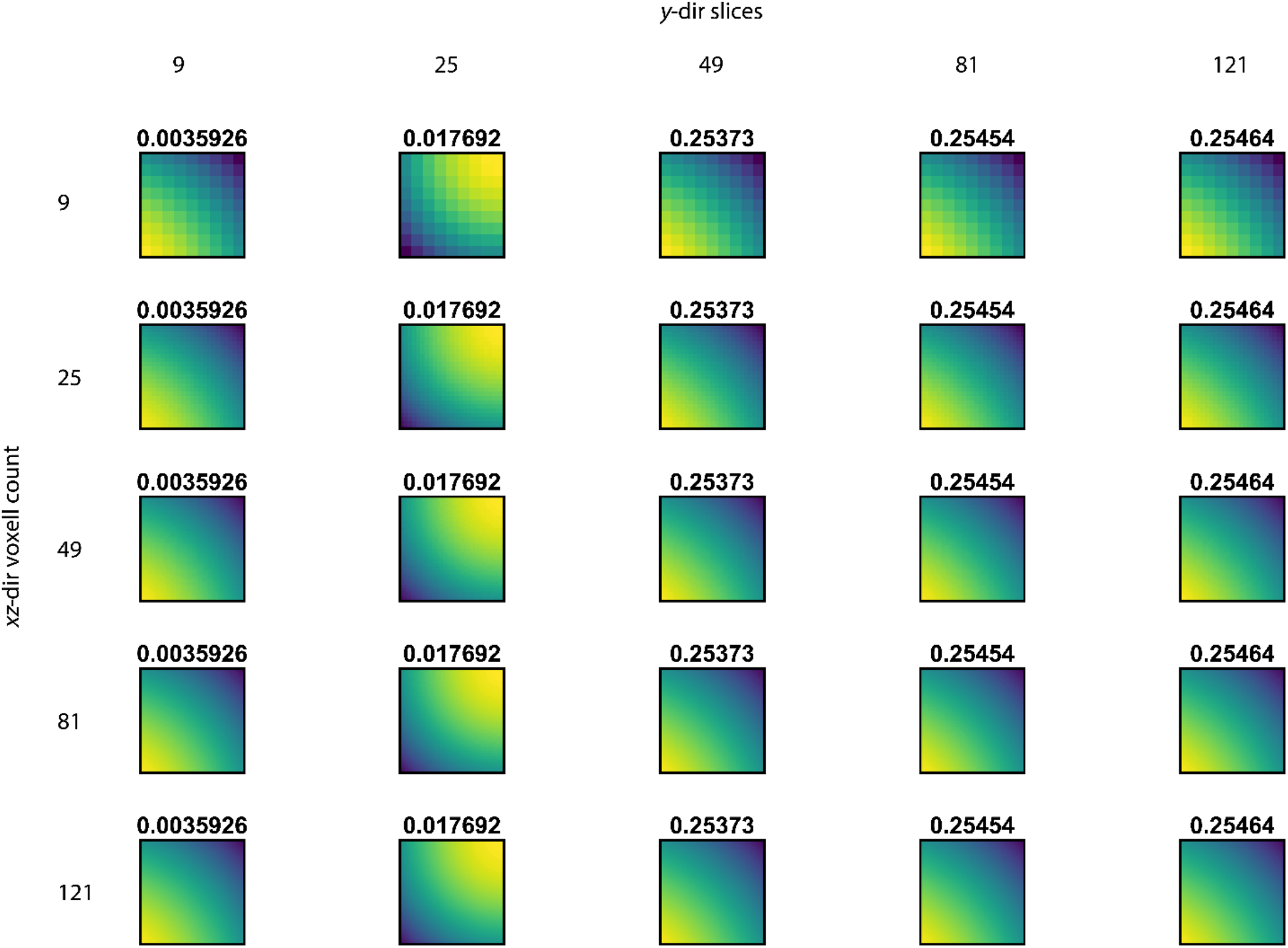
Similar to the previous figure, the values for the fundamental scattering model converge at around 121 *y*-slices to the reported values in the main text. Changing the voxel count in the *xz* plane does not affect the reported phase shift.

